# Adrenal remodeling encodes stress axis memory to shape fentanyl seeking

**DOI:** 10.64898/2026.10.20.760432

**Authors:** P. Maretich, T. M. Cannon, T. Milo, N. Pagane, E. J. Frey, F. Choe, E. Vargas Paniagua, R. Leomi, S. Machen, K. Nagao, H. S. Wong, S. L. Prescott, K. J. Ressler, P. Anikeeva

## Abstract

The hypothalamic-pituitary-adrenal (HPA) axis is implicated in substance use disorders, yet the features of this circuit that contribute to addiction vulnerability remain unclear. Here, we show that fentanyl engages with different components of the HPA axis, and this engagement varies across timescales. Within minutes, fentanyl activated corticotropin-releasing hormone neurons in the paraventricular nucleus of the hypothalamus (PVN^CRH^) and transiently increased adrenocorticotropic hormone (ACTH) levels. Hours to days later, fentanyl suppressed PVN activity, reduced adrenal size, and impaired corticosterone release. Conversely, ACTH-induced adrenal hypertrophy, increased corticosterone, suppressed fentanyl-evoked PVN^CRH^ activity, and elevated fentanyl-seeking behavior. These findings identify adrenal remodeling as a physiological record of stress history that shapes neuroendocrine and behavioral responses to fentanyl, mechanistically linking stress and addiction while revealing treatment targets for opioid dependence.

---

Despite widespread opioid exposure, only a subset of those exposed develop opioid use disorder (*1*). A meta-analysis of newly incident heroin users found that 23-38% developed dependence within the first year of use, consistent with evidence that addiction risk varies across individuals (*2–5*). Stress physiology represents one candidate source of this variability: both excessive and blunted responses to stressors have been associated with addiction risk in humans and model organisms (*6–8*). The hypothalamic-pituitary-adrenal (HPA) axis is an evolutionarily conserved neuroendocrine pathway that coordinates physiological stress responses and has been implicated in addiction vulnerability and drug seeking (*9–12*). Within this circuit, corticotropin-releasing hormone (CRH)-producing neurons (PVN^CRH^) in the paraventricular nucleus of the hypothalamus (PVN) drive pituitary adrenocorticotropic hormone (ACTH) release and downstream adrenal glucocorticoid production. Prolonged HPA axis activity can alter adrenal gland mass and hormone output (*13, 14*). Clinical studies indicate that opioids perturb this system; opioid-induced adrenal insufficiency occurs in 9–29% of patients prescribed opioids long-term, and regular heroin users exhibit abnormal daily cortisol rhythms (*8, 15–19*). Together, these findings suggest that opioid exposure engages a dynamic stress-endocrine circuit which may shape individual vulnerability to compulsive drug seeking. However, the mechanisms by which opioids alter HPA axis tone, and whether this perturbation causally shapes downstream neural, endocrine, and behavioral responses, remain unclear.

To address these open questions, we examined how the synthetic opioid fentanyl, involved in the majority of overdose deaths, engages different components of the HPA axis over time (*20*). We combined cell type-specific PVN^CRH^ recordings, electrophysiology, hormone measurements, multisite neural recordings, adrenal ultrasound, RNA sequencing, and pharmacological manipulation to characterize HPA axis dynamics across hypothalamic, adrenal, and mesolimbic circuits. Further, we predicted that ACTH-driven adrenal remodeling would amplify drug-seeking behavior. To test this causal relationship, we measured fentanyl self-administration and physiological and behavioral responses to fentanyl in mice pretreated with ACTH.

## Fentanyl acutely activates the HPA axis

We first characterized the acute HPA axis response to a single dose of fentanyl (Fig. 1A). Contrary to the canonical sedative properties of opiates, in the minutes following fentanyl administration (0.4 mg·kg^−1^, subcutaneous, s.c.) we observed activation in CRH neurons within the PVN (PVN^CRH^) using fiber photometry in Crh-Cre driver mice virally transduced to express calcium indicator GCaMP6s in a Cre-dependent manner (Fig, 1B, C, Methods) (*21*). This activation was blunted by the pretreatment with the μ-opioid receptor antagonist, naloxone hydrochloride (5 mg·kg^−1^, s.c.), 5 min prior to fentanyl injection (Fig. 1D). Additionally, fentanyl rapidly produced the Straub response, a rigid curling of the tail toward the head associated with μ-opioid receptor activation, accompanied by frequency-specific changes in PVN local field potential (LFP) spectral power (Fig. 1E) (*22, 23*). Straub epochs were associated with significantly reduced delta (δ), theta (θ), and alpha (α) power and a significantly increased high-gamma (γ) power (Fig. 1F). Furthermore, significant upregulation of the neuronal activity marker, c-Fos, was observed in the PVN regions of mice (wild type, WT) treated with fentanyl (0.4 mg·kg^−1^, intraperitoneal, i.p.) but not with saline (Fig. 1G, H).

**Fig. 1.**
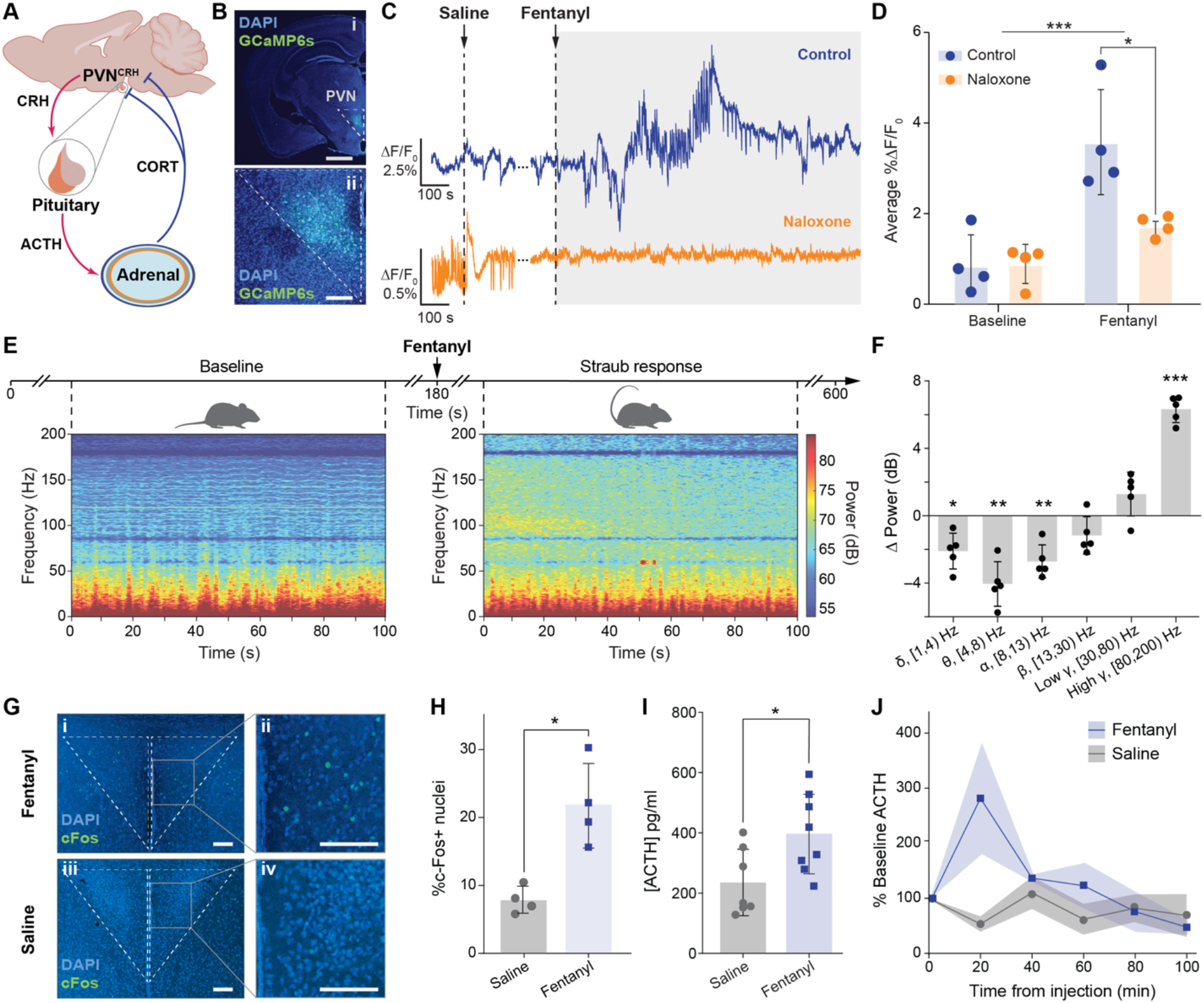
Acute fentanyl exposure rapidly activates PVN^CRH^ neurons. **(A)** HPA axis schematic. **(B)** Coronal brain slice of Crh-Cre mouse expressing AAV9-hSyn::FLEx-GCaMP6s (FLEX::GCaMP6s, ≥ 1×10^13^ vg/mL). Scale bars = 500 µm (top), 100 µm (bottom) **(C)** Representative traces of normalized GCaMP6s fluorescence (ΔF/F_0_) in CRH^PVN^ neurons at baseline (top) and pretreated with naloxone (bottom) in response to saline and fentanyl. **(D)** Mean normalized fluorescence change ΔF/F_0_ for fentanyl or saline treated mice with or without naloxone pretreatment. Two-way ANOVA: main effect of fentanyl, F(1,12) = 24.14, P = 0.0004; naloxone × fentanyl interaction, F(1,12) = 6.790, P = 0.0230 (n = 4). **(E)** Group-averaged, log-transformed, PVN LFP spectrograms (n = 5). **(F)** Band power difference between baseline and Straub epochs. Repeated-measures ANOVA showed Band × Epoch interaction (Greenhouse–Geisser p = 1.84 × 10^-6^); FDR-corrected paired t-tests showed decreased delta (p = 0.0179), theta (p = 0.0071), and alpha (p = 0.0071), increased high gamma (p = 0.00032) (n = 5). **(G)** Confocal micrographs of coronal PVN sections immunostained for c-Fos from mice treated with fentanyl (top) or saline (bottom). Scale bars = 100 µm. **(H)** Percentage of c-Fos expressing nuclei in the PVN of fentanyl and saline treated mice. Welch’s t-test, t(3.606) = 4.296, p = 0.0158 (n = 4). **(I)** Circulating ACTH following saline (n = 7) or fentanyl (n = 8) injections. Welch’s t-test, t(12.99) = 2.610, p = 0.0216. **(D, F, H, I)** Bars and error bars represent mean ± standard deviation (SD). **(J)** ACTH normalized to baseline after saline or fentanyl injections. Two-way ANOVA: treatment effect F(1,24) = 4.775, p = 0.0389; treatment × time interaction F(5,24) = 2.976, p = 0.0314. Lines and shaded areas represent mean ± standard error of the mean (SEM).

Consistent with broader HPA axis activation, plasma samples taken 20-30 min after saline or fentanyl administration showed increased circulating ACTH in the latter group (397.3 ± 132.1 pg/ml) relative to the former (233.6 ± 110.8 pg/ml, Fig. 1I). Prior studies showing rodent ACTH hypersecretion in response to opioids indicated that the effects peak within 30 min (*24*). Similarly, repeated plasma sampling from jugular-catheterized mice showed a transient ACTH increase 20 min after fentanyl administration that returned to baseline within 100 min (Fig. IJ) (*25, 26*). Interestingly, ACTH shares a common precursor molecule with the endogenous opioid peptide β-endorphin; subsequently, the two are generally released in equimolar ratios. We found that fentanyl treatment did not alter circulating levels of β-endorphin, suggesting that their relative abundance can vary depending on the conditions under which proopiomelanocortin (POMC)-derived peptides are processed and secreted (*27–29*) (Fig. S1).

## Activation of PVN^CRH^ neurons following fentanyl exposure is accompanied by dopamine release in the reward circuit

PVN^CRH^ neurons send direct projections to the ventral tegmental area (VTA), a node of the mesolimbic reward pathway enriched in dopaminergic neurons (*30–34*). Consequently, we hypothesized that the acute hormonal response may coordinate with neural activity in reward-processing circuits during fentanyl exposure. To test this, we expressed Cre-dependent GCaMP6s in PVN^CRH^ neurons, relying on anterograde transport for VTA fluorescence, and broadly expressed dopamine sensor dLight1.3b in the nucleus accumbens (NAc), which receives dopaminergic projections from the VTA (Fig. 2A, B). A custom multi-fiber imaging device then enabled simultaneous recording of activity in the PVN, VTA, and the NAc (Fig. 2C). Within this probe, two flexible optical multicore fibers each containing 7400 independent 4 µm waveguides provided spatially resolved GCaMP6s imaging in the PVN and VTA and a single-core polymer fiber in the NAc captured bulk dLight1.3b activity.

**Fig. 2.**
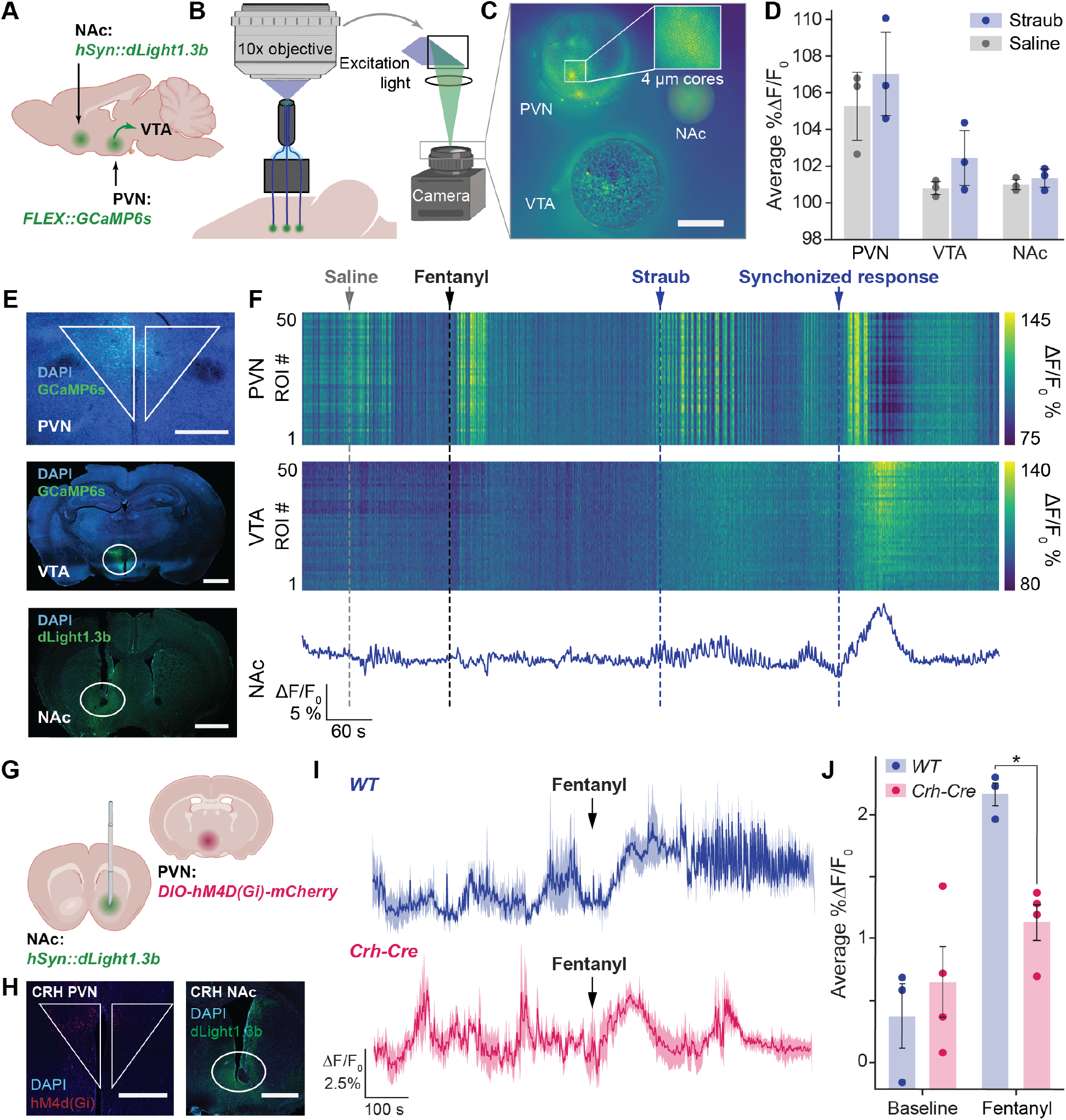
Multi-site neural imaging with spatial resolution links stress and reward circuits. **(A)** Injection of AAV9-hSyn::dLight1.3b into the NAc and AAV9-FLEX::GCaMP6s into the PVN. An arrow represents anterograde labeling from PVN^CRH^ to VTA. **(B)** Multisite probe for spatially resolved imaging in the PVN and VTA and NAc photometry deployed with a 10X objective lens and a camera operating at 10 fps. **(C)** Camera field of view. Scale bar = 200 µm. **(D)** Normalized GCaMP6s (PVN and VTA) and dLight1.3b (NAc) ΔF/F_0_ during saline and Straub epochs. Bars and error bars represent mean ± S.D (n = 3). **(E)** Confocal micrographs corroborate expression of GCaMP6s in the PVN^CRH^ cell bodies and in PVN^CRH^→VTA projections, and dLight expression in the NAc. Scale bars = 500 µm. **(F)** Simultaneous imaging of putative PVN^CRH^ neurons GCaMP6s ROIs (top), their terminals in the VTA (middle), and dLight1.3b fluorescence in the NAc (bottom) showing fentanyl-associated increases in PVN^CRH^ activity and NAc dopamine release during Straub response. **(G, H)** Expression of hM4D(Gi)-mCherry in the PVN^CRH^ and dLight1.3b in the NAc in Crh-Cre mice. Scale bars = 500 µm. **(I)** dLight1.3b fluorescence (ΔF/F_0_) in the NAc in the Crh-Cre and WT mice following pretreatment with CNO and fentanyl administration. Lines and shaded areas represent mean ± SEM. **(J)** Two-way repeated measures ANOVA revealed a significant main effect of treatment (Baseline vs Fentanyl: F(1,5) = 50.38, p = 0.0009) and a significant treatment × genotype interaction (F(1,5) = 15.41, p = 0.0111), indicating differential fentanyl sensitivity between genotypes. Bars and error bars represent mean ± SD. Crh-Cre n = 4; WT n = 3.

Robust coordinated activity of putative PVN^CRH^ neurons (regions of interest (ROI) within the multicore fiber images, Methods) and their terminals in the VTA was observed in response to fentanyl injections (0.4 mg·kg^−1^, s.c.) and during the Straub response (Fig. 2D, E). Opioids characteristically generate short bursts of dopamine (*35*). These bursts were observed during the Straub period, and were synchronous with PVN^CRH^ neuron activity (Fig. 2F).

To elucidate whether PVN^CRH^ activity functionally contributes to fentanyl-evoked dopamine release in the NAc, an inhibitory designer receptor hM4d(Gi) was virally expressed in a Cre-dependent manner in Crh-Cre (n = 4) or WT mice (n = 3). In the same subjects, dLight1.3b was broadly expressed in the ipsilateral NAc to enable photometric readout through a single-core optical fiber (Fig. 2G, H). Pre-treatment with clozapine N-oxide (CNO; 10 mg·kg^−1^, s.c.) to trigger hM4d(Gi) 10 min prior to fentanyl administration resulted in significantly blunted dopamine release in the minutes (250-450 s) following fentanyl administration in Crh-Cre mice as compared to WT controls, indicating that PVN^CRH^ activity contributes to fentanyl-induced reward (Fig. 2I, J) (*36, 37*).

## At longer timescales fentanyl exposure inhibits PVN activity, induces adrenal atrophy, and blunts corticosterone release

To reconcile the well-established sedative properties of fentanyl with our observation of acute PVN^CRH^ neuron activation, we tested whether the effects of fentanyl on the brain change in the hours following administration (*38*). Within 5 hours following fentanyl injection, in contrast to the acute effects, electrophysiology recordings in the PVN revealed significantly lower average LFP power in fentanyl-injected animals compared to saline controls (Fig. 3A). The reduction was not specific to any individual frequency band between 1-200 Hz (Fig. 3B, S2). These data suggest that fentanyl leads to extended suppression of PVN activity over time.

**Fig. 3.**
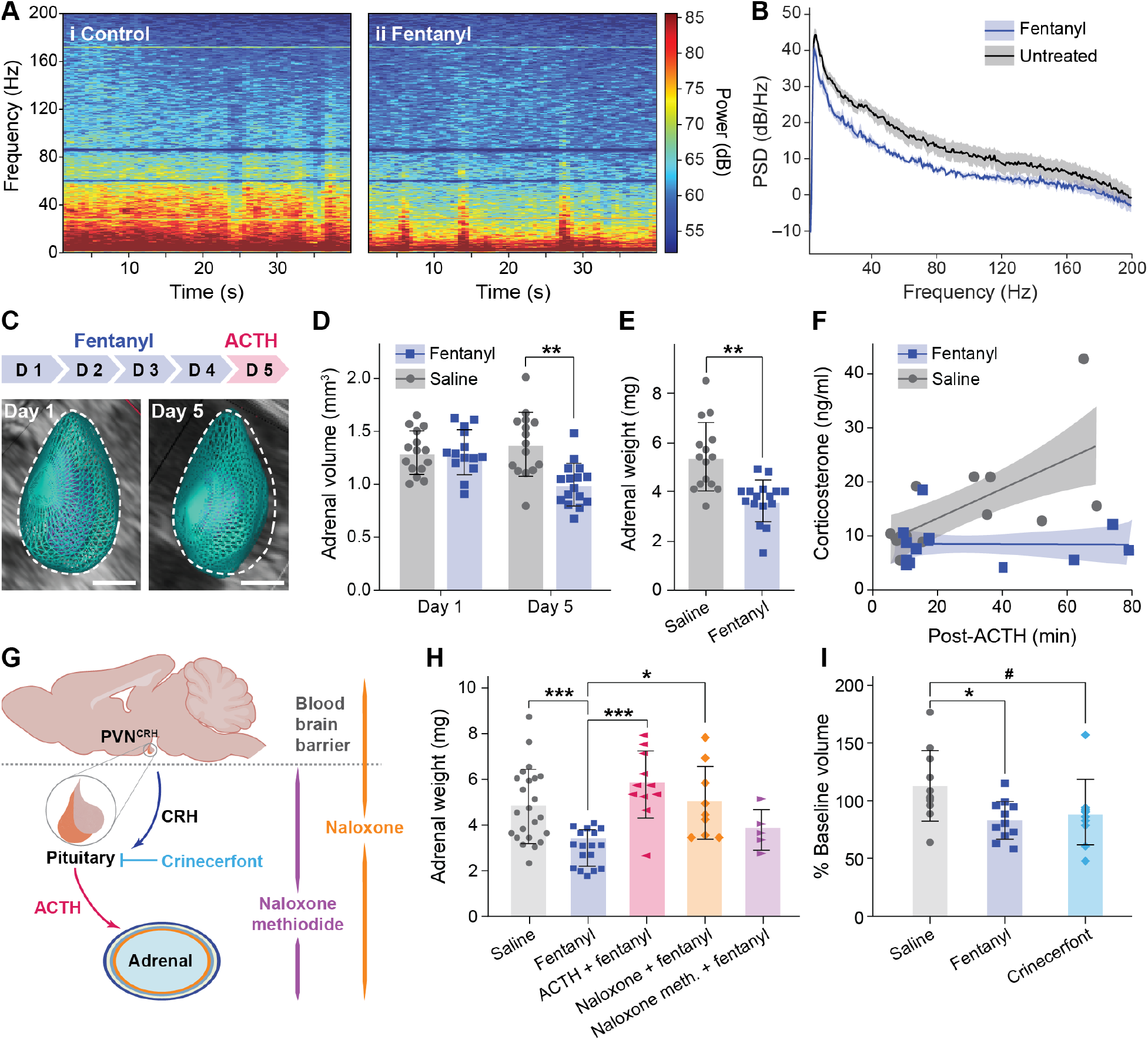
Fentanyl acts centrally to induce adrenal atrophy over hours and repeated doses. **(A)** Group-averaged PVN LFP spectrograms 5 h post treatment (n = 3 and n = 4 for treated and untreated mice, respectively) with a shared power scale. **(B)** Frequency-resolved LFP power spectral density for fentanyl-treated and untreated mice. Lines and shaded areas represent mean ± SEM. **(C)** Ultrasound-derived adrenal volume reconstruction at baseline (left) and following 4 days of fentanyl administration within the same adrenal gland. Scale bar = 1 mm. **(D)** Adrenal volume recorded within the same fentanyl- or saline-treated subjects on days 1 and 5. Two-way repeated measures ANOVA revealed a significant treatment × time interaction (F(1,14) = 19.32, p = 0.0006). Šídák’s post-hoc comparisons: day 1 fentanyl vs. saline, p = 0.9998 (ns); day 5 fentanyl vs. saline, p = 0.0017; fentanyl day 1 vs. day 5, p = 0.0003; saline day 1 vs. day 5, p = 0.4929 (ns). n = 8 per group. **(E)** Adrenal mass on day 5 for fentanyl and saline treated mice. Mann-Whitney, U = 4, p = 0.0019. n = 8 per group. Bars and error bars represent mean ± SD. **(F)** Corticosterone concentration plotted as a function of each mouse’s blood collection time relative to ACTH injection. Simple linear regression, 95% confidence intervals; saline: Y = 0.272X + 8.77; F_(1_,_22)_ = 15.31, p = 0.0007, R^2^ = 0.41; fentanyl: Y = −0.002X + 9.42; F_(1_,_23)_ = 0.005, p = 0.94, R^2^ = 0.0002; n = 13 per group. Lines and shaded areas represent mean ± SEM. **(G)** Pharmacological manipulation across the HPA axis. **(H)** Adrenal weight in mice treated for 4 days with saline (n = 12), fentanyl alone (n = 10), fentanyl and ACTH (n = 6), fentanyl and naloxone (n = 5), and fentanyl and naloxone methiodide (n = 3). Two-way ANOVA revealed a significant main effect of treatment (F(4,20) = 8.416, p = 0.0004), indicating differential effects across intervention groups. Dunnett’s post-hoc comparisons with fentanyl as the reference control indicated significant differences between fentanyl alone and saline (p = 0.0004), fentanyl alone and fentanyl + ACTH (p = 0.0008), fentanyl alone and fentanyl + naloxone (p = 0.0411). **(I)** Change in adrenal volume on day 5 with respect to day 1 baseline in mice treated with saline (n = 6), fentanyl (n = 6), and crinecerfont (n = 5). Two-way ANOVA revealed a significant main effect of treatment (F (2, 25) = 4.733, p = 0.0181). Tukey’s post-hoc comparisons showed fentanyl significantly reduced adrenal volume change relative to saline (p = 0.0192); crinecerfont vs. saline (p = 0.0774); fentanyl vs. crinecerfont (p = 0.893). **(D, E, F, H, I)** Bars and error bars represent mean ± SD.

Given the central role of PVN^CRH^ neurons in regulating the HPA axis, we hypothesized that this repeated suppression of PVN activity may have downstream consequences in the periphery, namely the adrenal glands, that outlast fentanyl exposure (*39*–*41*). We therefore investigated the effects of repeated fentanyl exposure on adrenal morphology and function. Six-week-old WT mice received daily fentanyl (0.3 mg·kg^−1^, i.p.) or saline injections for 4 days, and their adrenal volume was measured via ultrasound prior to (day 1) or following (day 5) final administration. At day 5, we additionally conducted an ACTH stimulation test (1 mg·kg^−1^, i.p.) to assess adrenal responsiveness (Fig. 3C). Volumetric reconstructions of the same adrenal glands over time revealed a ~ 25% decrease in volume corresponding to a ~1.8 mg decrease in weight of the adrenal glands from fentanyl-treated mice as compared to saline controls by day 5 (Fig. 3D, E). Final ultrasound-derived adrenal volumes correlated with dissected adrenal weights collected on the same day (R^2^ = 0.72, Fig. S3A), and no significant differences in brain or body weight were observed in fentanyl-treated animals as compared to controls (Fig. S3B, C). Linear regression analysis of the ACTH challenge revealed a significant positive relationship between circulating corticosterone minutes following pharmacological stimulation in saline controls, but not in fentanyl-exposed subjects (Fig. 3F) (*42*). Together, these findings indicate that repeated fentanyl exposure is sufficient to reduce adrenal size and blunt corticosterone release.

To further characterize the effects of fentanyl on adrenal tissue, we performed bulk RNA sequencing on adrenal glands from mice treated with fentanyl (0.3 mg·kg^−1^, i.p.) or saline for 4 days. Across treatment conditions, inference showed that approximately 75% of the bulk adrenal transcriptomic signal derived from cells in the zona fasciculata (ZF) (Fig. S4A), the primary source of corticosterone. Gene-concept network plots showed enrichment of collagen- and extracellular matrix-associated programs driven by genes including *Itga2, F2r*, and *Prkcd*, while downregulated genes including *Mtor* and *Abcd1* were associated with pathways related to cellular growth, metabolism, and adrenocortical function (Fig. S4B, C). Consistent with these patterns, adrenal glands from fentanyl-treated mice showed enrichment of collagen biosynthesis and stress-response pathways, together with decreased representation of transcriptional, translational, and metabolic programs (Fig. S5; Table S1).

The observed functional and transcriptional changes suggested that fentanyl exposure reduces adrenal trophic support but did not causally establish where in the body fentanyl acts to produce this remodeling. Since μ-opioid receptors are broadly distributed, we tested whether fentanyl-induced adrenal atrophy could arise from opioid receptor signaling in the brain, pituitary, adrenal gland, or peripheral tissues (*43*). To distinguish central from peripheral signaling, we co-administered fentanyl with either BBB-permeable naloxone hydrochloride or BBB-impermeable naloxone methiodide (Fig. 3G). Co-administration with naloxone hydrochloride rescued adrenal weight to values not significantly different from saline controls. In contrast, co-administration with naloxone methiodide did not significantly increase adrenal weight in fentanyl-treated animals (Fig. 3H). Notably, administration of naloxone hydrochloride or naloxone methiodide alone did not lead to changes in adrenal weight (Fig. S6; Table S2). These experiments indicated that fentanyl acts centrally, likely on PVN^CRH^ neurons, to induce adrenal atrophy.

This result suggested that adrenal atrophy could be a direct result of reduced signaling downstream of PVN^CRH^ neurons activity, potentially from reductions in ACTH levels. To test whether ZF activity and volume are dynamically regulated by ACTH, we co-administered this hormone during the 4-day fentanyl exposure paradigm (*44, 45*). Unlike the subjects treated with fentanyl alone, whose adrenal volume was significantly reduced, the mice co-treated with ACTH exhibited adrenal volumes not significantly different from the saline-treated controls (Fig. 3H, Table S2). This suggested that fentanyl-induced reduction of ACTH is responsible for adrenal remodeling, and that restoring adrenal trophic support is sufficient to prevent adrenal tissue loss associated with fentanyl exposure.

Since CRH receptor signaling drives ACTH release and subsequent adrenal function, we tested whether blocking this pathway is sufficient to produce adrenal atrophy. Subsequently mice were treated with crinecerfont, a CRH type 1 receptor antagonist, and their adrenal volume was recorded by ultrasound (*46*). Following a 3-day treatment with crinecerfont, mice showed adrenal volume loss (87 ± 19% of baseline, n = 6) comparable to fentanyl (82 ± 16%, n = 6), though this did not reach statistical significance versus saline controls (p = 0.0774; Fig. 3I). These data provide further evidence that fentanyl-induced adrenal atrophy is at least, in part, driven by disruption in CRH-to-ACTH signaling within the HPA axis.

To determine whether adrenal morphological changes reflect altered receptor expression rather than the sustained suppression of PVN activity, we quantified HPA axis-associated transcripts (*Mc2r, Mrap, Nr3c1, Fkbp5, Pomc, Crhr1, Crhr2*, and *Crhbp*) in bulk adrenal tissue from fentanyl-treated and saline control mice (Fig. S7). Across these transcripts, treatment had no significant effect on normalized expression (F(1,176) = 0.276, p = 0.60) (*47*). Together, these results are consistent with fentanyl-induced suppression of the PVN contributing to adrenal tissue remodeling in lieu of alterations in adrenal receptor expression.

## Elevated ACTH alters fentanyl-induced PVN activity and promotes fentanyl seeking

Adrenal remodeling reflects sustained deviations in HPA axis activity, which raises the question of how adrenal hormones feed back to alter the brain’s response to fentanyl and subsequent drug-seeking behaviors. Our data indicate that the rewarding properties of fentanyl are relayed, in part, through the HPA axis. This link prompted us to investigate the mechanism underlying the comorbidity of stress and opioid addiction (*48*). Chronic stress reshapes the HPA axis’s structural and functional capacity through adrenal remodeling, a process that unfolds over hours to days and cannot be resolved by snapshot hormone measurements. Our data showing fentanyl’s multiscale engagement with the HPA axis underscored a need for a model capable of predicting how stress-induced changes in adrenal function influence drug responses. We therefore employed a dynamic mathematical model of HPA axis regulation, parameterized with literature-validated rates to ensure biological feasibility, to investigate the relationship between sustained stress and physiological responses to fentanyl. In this model, stress input (s) drives PVN^CRH^ activity, CRH stimulates ACTH release, and ACTH promotes adrenal corticosterone production, with the latter providing negative feedback onto upstream CRH and ACTH signaling (Fig. S8) (*49–55*). The model also includes adrenal functional mass as a slower state variable that changes according to ACTH-dependent cell-mass production and adrenal cell-mass removal (Table S3). Using this framework, we simulated 4 days of elevated ACTH, mimicking the hormonal changes associated with chronic stress. Relative to controls, adrenal functional mass was predicted to increase over a 4-day pre-treatment period, along with corticosterone output. Furthermore, the model predicted reduced CRH activity both at baseline and in response to fentanyl following ACTH-induced hypertrophy (Fig. 4A-C).

**Fig. 4.**
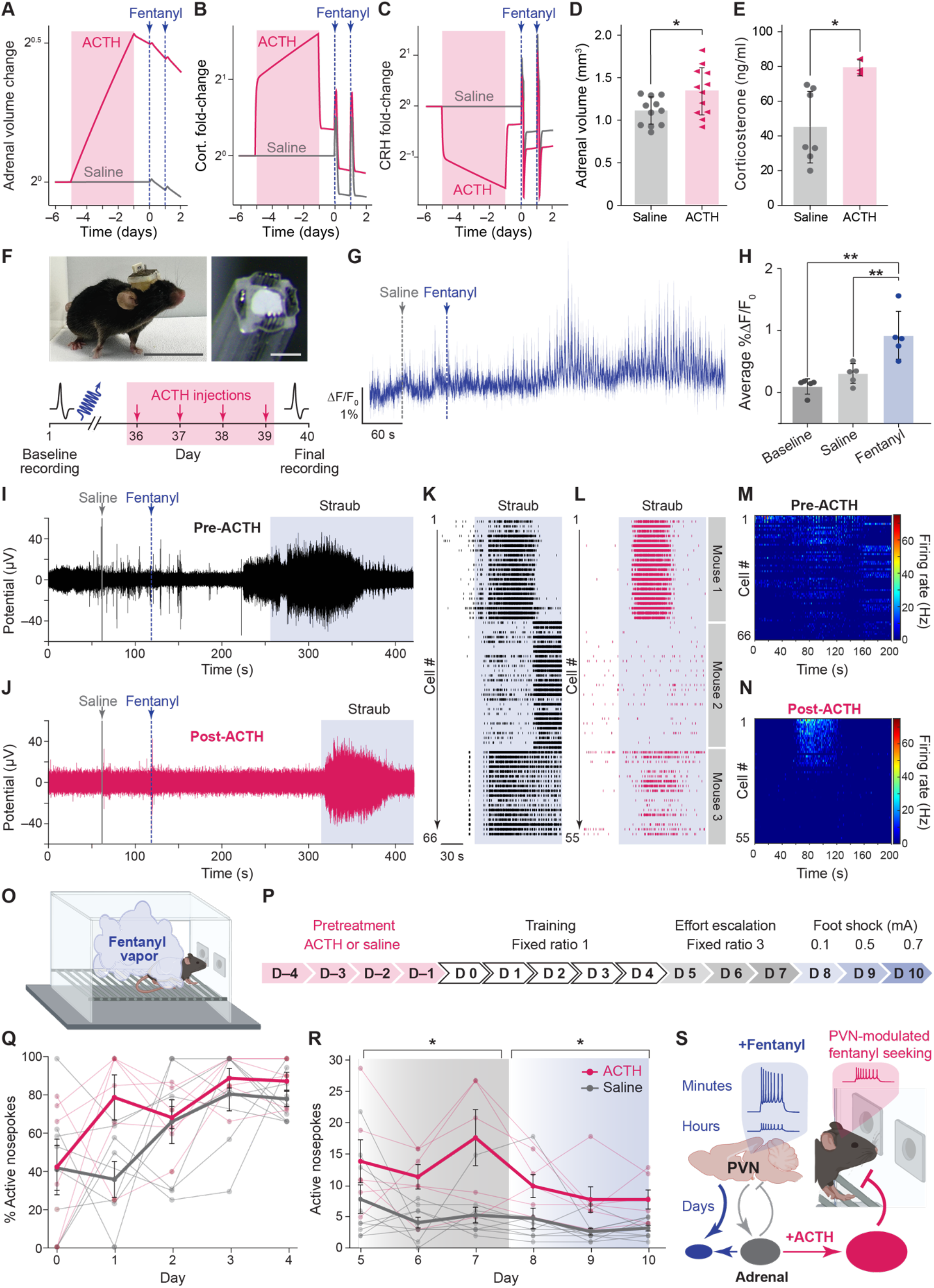
ACTH-induced adrenal growth alters physiological responses to fentanyl. **(A–C)** Mathematical modeling predicts that 4 days of elevated ACTH input increases adrenal functional mass and corticosterone output while suppressing CRH activity both at baseline and in response to fentanyl. **(D)** ACTH pretreatment (0.45 mg·kg^−1^, i.p., daily × 4 days) significantly increased adrenal volume measured by ultrasound (ACTH: 1.321 ± 0.278 mm^3^, n = 12 vs. Saline: 1.089 ± 0.163 mm^3^, n = 11; Mann-Whitney, U = 33, p = 0.0424). **(E)** Plasma corticosterone increased in ACTH-treated mice (83.66 ± 5.42 ng/mL, n = 3) compared to saline controls (46.53 ± 19.24 ng/mL, n = 7; Mann-Whitney, U = 0, p = 0.0167). **(F)** Mouse post-implant with scale bar = 5 cm (upper left). Cross section of multimodal neural probe (12 tungsten electrodes + 300 μm optical waveguide) implanted in PVN for simultaneous electrophysiology and fiber photometry with scale bar = 300 μm (upper right). Experimental timeline shown below. **(G)** Group-level photometry signal showing mean %ΔF/F_0_ ± SEM (n = 5). **(H)** Quantification of %ΔF/F_0_ revealed significant increases in neural activity following fentanyl administration (baseline: 0.1052 ± 0.1286; saline: 0.2933 ± 0.1658; post-fentanyl: 0.9153 ± 0.3911; n = 5). Two-way ANOVA detected a significant effect of condition (F(2,8) = 17.10, p = 0.0013). Šídák’s post-hoc comparisons showed post-fentanyl activity significantly higher than baseline (p = 0.0010) and saline (p = 0.0053). **(I, J)** Single electrode spikeband at baseline (black) and post-ACTH (red). **(K, L)** Single-unit recordings from 66 neurons (3 mice) reveal elevated activity during Straub tail reflex pre-ACTH, with suppressed activity post-ACTH. **(M, N)** Heatmap of individual neural firing rate (Hz) at baseline (top) and post-ACTH (bottom). **(O, P)** Schematic of fentanyl vapor self-administration chamber enabling non-surgical operant behavior and experimental paradigm. **(Q)** Training phase (FR1, Days 0–4): Both groups learned to discriminate the active nosepoke, achieving >60% selectivity by Day 4. Mean ± SEM plotted in bold with individual mouse trajectories shown. **(R)** During increased effort (FR3: three nosepokes per dose; Days 5–7) and escalating electrical stimulation (FS: Days 8–10), ACTH-pretreated mice (red) showed significantly elevated active responding compared to saline controls (gray). FR3: ACTH 13.97 ± 5.85 vs. Saline 5.78 ± 2.31 nosepokes (p = 0.024*, +142%); FS: ACTH 8.36 ± 3.39 vs. Saline 3.56 ± 1.99 (p = 0.023*, +135%). Two-way ANOVA confirmed highly significant main effect of Group (F(1,76) = 35.20, p < 0.0001), significant Day effect (F(5,76) = 3.13, p = 0.013), and no Group × Day interaction (F(5,76) = 1.04, p = 0.398), indicating sustained ACTH-driven enhancement across testing. **(S)** Schematic summary: Fentanyl induces acute HPA axis activation, but long-term suppression and adrenal atrophy; ACTH-induced adrenal hypertrophy increases corticosterone output while suppressing PVN^CRH^ activity, resulting in sustained elevation of fentanyl-seeking behavior. **(D, E, H, N, R)** Bars and error bars represent mean ± SD. * p < 0.025 (Bonferroni-corrected for phase-level comparisons).

We experimentally tested this prediction in mice by administering 4 daily doses of either saline or ACTH (0.45 mg·kg^−1^, i.p.) and recording adrenal volume via ultrasound. ACTH-treated mice developed ~20% larger adrenal glands relative to saline controls (Fig. 4D). These results are consistent with previous findings demonstrating adrenal hypertrophy following 14 days of ACTH treatment (*56*). Consistent with our prediction that increased adrenal functional mass would enhance steroidogenic output, plasma corticosterone was also increased in ACTH-treated mice relative to saline controls (Fig. 4E). Taken together, the adrenal gland behaves as a cumulative record of HPA axis history: repeated fentanyl exposure reduced adrenal size and corticosterone output, whereas ACTH pretreatment enlarged the adrenal gland and increased baseline corticosterone.

To interrogate how adrenal hypertrophy shapes fentanyl-evoked PVN^CRH^ responses, we deployed custom multifunctional fiber-based probes comprising 12 tungsten electrodes (20 µm diameter) arranged around a 300 µm diameter polymer optical waveguide in mice treated with ACTH and fentanyl. These devices were applied in Crh-Cre mice virally transduced to express GCaMP6s in PVN^CRH^ neurons (Fig. 4F). The majority of PVN neurons express CRH (~80%), nevertheless photometric recordings were used to corroborate that high-resolution neural activity concomitantly recorded via electrodes reflected HPA-axis electrophysiology (*57*). As shown, a significant increase in GCaMP6s % ΔF/F_0_ in the PVN^CRH^ population was observed following acute fentanyl administration (Fig. 4G, H). In the same subjects, electrophysiology recordings revealed fentanyl-evoked increases in PVN neuron firing, which coincided with the Straub tail reflex (Fig. 4I). In contrast, although the spike waveforms remained stable, recordings from the same electrodes following the 4-day ACTH treatment revealed altered activity in response to fentanyl (Fig. 4J, S9A–C). Single-unit recordings from 66 neurons (n = 3 mice) showed increased activity during Straub, which was delayed (104.2 ± 85.01 s) or suppressed following ACTH treatment (Fig. 4K-N, S9D). Within 4 days, ACTH-induced adrenal changes were paired with altered PVN^CRH^ responses to fentanyl, suggesting that adrenal state may shift on a timescale relevant to early drug exposure.

We then hypothesized that these changes in fentanyl-evoked PVN^CRH^ activity would have consequences for drug-seeking behavior in mice with ACTH-induced adrenal hypertrophy. To avoid the confounds of opioid analgesic effects and minimize handling-associated stress (*58–60*), we adapted a non-surgical vapor-based self-administration paradigm via a custom chamber equipped with two nose-poke ports and a floor grid capable of delivering current (Fig. 4O, Fig. S10A–D, Table S4). Mice were pre-treated with ACTH (or saline, co-housed littermates) for 4 days as described above (days –4 to –1) and subjected to a fentanyl self-administration paradigm consisting of 5 days of training (days 0 to 4), followed by effort escalation (days 5–7), and finally increasing foot-shock punishment (days 8–10) (Fig. 4P). During the training phase, mice learned to discriminate the active nose-poke, reaching >60% active responses by day 5, with ACTH- and saline-treated animals learning at similar rates (Fig. 4Q). When effort was increased to 3 active nose-pokes per dose of fentanyl (FR3), ACTH-treated mice performed 142% more active nose-pokes than saline controls. When fentanyl delivery was paired with escalating foot shock (FS), ACTH-treated mice sustained significantly elevated active responses compared to saline controls. A two-way ANOVA revealed that ACTH-treated mice showed significantly elevated active responses compared to saline controls across days 5–10 (F(1,76) = 35.20, p < 0.0001). There was also a significant main effect of Day (F(5,76) = 3.13, p = 0.013), indicating that behavior evolved over time. The Group × Day interaction was not significant (F(5,76) = 1.04, p = 0.398), demonstrating that the effect of ACTH pretreatment did not differ significantly across days (Fig. 4R).

These findings support a link between ACTH-induced HPA axis remodeling and increased fentanyl self-administration. Fentanyl acutely activates PVN^CRH^ neurons contributing to VTA activation and dopamine release in the NAc (Fig. 2). This acute activation is followed by sustained PVN^CRH^ suppression, which contributes to adrenal atrophy (Fig. 3). In contrast, elevated ACTH induces adrenal hypertrophy manifesting in elevated corticosterone and suppressed PVN^CRH^ responsiveness to fentanyl, which is accompanied by heightened drug-seeking behavior under conditions of increasing cost and punishment (Fig. 4S).

## Discussion

By combining physiological recordings of neural activity in the PVN, endocrine and ultrasound characterization of the adrenals, mathematical modeling, and behavioral assays, we show that the adrenal gland state is not merely a consequence of HPA axis activity but may also be a determinant of subsequent neuroendocrine, neural, and behavioral responses to fentanyl.

ACTH-induced adrenal hypertrophy altered fentanyl seeking, suggesting that prior HPA axis activation can shape later drug-directed behavior. One interpretation is that adrenal hypertrophy elevates baseline corticosterone, which may modify subsequent fentanyl-evoked HPA responses and reward circuit sensitivity. Our neurophysiology recordings provide a potential link between adrenal state and reward circuitry: chemogenetic inhibition of PVN^CRH^ neurons attenuated fentanyl-evoked dopamine release in the NAc, while ACTH-induced adrenal hypertrophy delayed and blunted fentanyl-evoked PVN^CRH^ neuron activity. Together, these findings suggest that adrenal state can alter the neural response to fentanyl, potentially changing the balance between reward and anti-reward dynamics driving compulsivity (*61*).

Although this study focused on the HPA axis, other stress systems, opioid-sensitive circuits, autonomic pathways, immune signals, and extra-hypothalamic brain regions are known to contribute to fentanyl responses and should be considered in addiction vulnerability (*61–64*). Their contribution to stress state-dependent changes in drug valence warrants further investigation. While PVN^CRH^ neurons participate in the HPA axis response to fentanyl, our data do not establish direct fentanyl action on these neurons. Fentanyl may instead alter PVN activity through upstream opioid-sensitive inputs or broader network effects (*65–68*). Indeed, the observed acute PVN^CRH^ activation in response to fentanyl may represents PVN^CRH^ disinhibition Finally, the exposure paradigms used in our study were modest relative to chronic stress or long-term opioid treatment to avoid compensatory changes or physical-dependence confounds (*69–73*). ACTH treatment and altered glucocorticoid tone target other tissues that could contribute to motivation (*74, 75*). Nevertheless, our findings identify adrenal state as a dynamic regulator of fentanyl response and suggest that recent stress can shape vulnerability to drug seeking. More broadly, they provide mechanistic insight into the comorbidity between stress-related disorders and addiction, highlighting how prior stress exposure can alter the brain’s response to drugs of abuse.

## Acknowledgments

This work was funded in part by the Director’s Pioneer Award from the National Institutes of Health and National Institute for Complementary and Integrative Health (DP1-AT011991), McGovern Institute for Brain Research at MIT, and Lore Harp McGovern Addiction Initiative at MIT. P.M. thanks support the Mathworks Fellowship and Janet and Sheldon Razin Fellowship. T.M.C. acknowledges the support of the NIH National Research Service Award (NRSA) F32 Fellowship. E.J.F. is a recipient of the National Science Foundation Graduate Research Fellowship (NSF GRFP). The custom neural interfaces deployed in this study were developed with the funding support of the K. Lisa Yang Brain-Body Center at MIT. The authors thank Gari Eberly for providing insightful edits and comments on the manuscript. The authors also acknowledge the use of the large language model Claude 4.5 Haiku (Anthropic, accessed January–September 2026) for grammatical assessment of sentences and refining data analysis scripts. All core intellectual content, scientific arguments, data interpretation, and original source code logic were conceived, written, and verified by the human authors, who maintain full accountability for the accuracy and integrity of the work. This work made use of the Preclinical Imaging & Testing Facility and BioMicro Center at MIT.

## Author contributions

Conceptualization: PM, PA

Methodology: PM, TMC, TM, NP, EJF, KN

Investigation: PM, TMC, TM, NP, EJF, KN, FC, EVP, SM

Visualization: PM, TMC, RL, PA

Funding acquisition: PA

Project administration: PA, RL, SM

Supervision: SLP, KJR, HSW, PA

Writing: All authors

## Data, code, and materials availability

All data supporting the findings of this study are available within the manuscript and supplementary information.

