## Supplementary materials for "Adrenal remodeling encodes stress axis memory to shape fentanyl seeking"

**The PDF file includes:**

Materials and Methods
Figs. S1 to S10
Tables S1 to S4
References cited only in the supplementary materials: 76 — 108

### Materials and Methods

#### Mice

All animal experiments were approved by the Massachusetts Institute of Technology (MIT) Committee on Animal Care (Protocol #2509000841). Mice were group-housed and kept on a 12-h light–dark cycle (07:00–19:00) with access to water and standard chow ad libitum. Mice undergoing self-administration assays were kept on a reverse light cycle (22:00–10:00). The facility maintained an ambient temperature of 18–23 °C and humidity of 40–60%. Male and female adult mice aged 6–30 weeks were used in all experiments excluding the following: male mice were used for the adrenal pharmacology rescue experiment, blood sampling via jugular catheters, and fentanyl self-administration assays. Female mice were used for the multisite imaging and photometry recordings. The following mouse strains were purchased or bred in-house and used directly: C57BL/6J (JAX 000664) and Crh-IRES-Cre (JAX 012704).

#### Ultrasound recordings and analysis

Adrenal ultrasound imaging was performed via Vevo 3100 Imaging System equipped with the LAZR-X Photoacoustic capability (Fujifilm VisualSonics) and a 70 MHz linear-array transducer to enable high-resolution visualization of adrenal structures. Mice were shaved over the abdominal region 24 h prior to imaging to optimize acoustic coupling. On the day of recording, animals were anesthetized with isoflurane (3% in oxygen) and positioned prone on a temperature-controlled heated platform to maintain physiological body temperature throughout the procedure. Ultrasound gel was applied to ensure proper transducer contact, and imaging was conducted under continuous anesthesia to minimize motion artifacts. Adrenal glands were identified using anatomical landmarks including the spine (medially), ipsilateral kidney (caudally), and rib cage (cranially), positioned at the cranial pole of each kidney in the retroperitoneal space. Adrenal glands were reconstructed as 3-dimensional (3D) structures, and volumes were calculated using the FujiFilm-VisualSonics VevoLabs software.

#### Adrenal dissection and weighing

Mice were euthanized by $CO_2$ exposure followed by cervical dislocation, and adrenal glands were dissected bilaterally under a stereomicroscope within 5 minutes. Surrounding fat and connective tissue were carefully removed before weighing. Left and right adrenal glands were weighed individually when both glands were available (Mettler Toledo, 30697411). Glands that were cut or damaged during dissection were excluded from the analysis.

#### Statistical analysis of adrenal volume and weight

Both left and right adrenals were plotted in all figures; however, each mouse served as the statistical unit (n=1) for all tests. Statistical significance was set at $\alpha = 0.05$. For single pairwise comparisons between two groups, the Mann-Whitney U test (nonparametric alternative to t-test) was used when data were non-normally distributed or sample sizes were small. Welch's t-test with unequal variance assumption was used for normally distributed data with unequal sample sizes (*76*). For experiments with multiple comparisons against a single control group, Dunnett's test was used to control family-wise error while maximizing statistical power. For experiments with multiple pairwise comparisons among all groups without a designated control, two-way ANOVA (ordinary or repeated measures as appropriate) was performed to test for overall treatment effects. When two-way ANOVA revealed significant main effects or interactions (p <

0.05), Tukey's multiple comparisons test was applied to compare multiple treatment combinations or timepoints (*77*). Repeated measures ANOVA was used when the same animals were measured across multiple timepoints or conditions. All statistical tests were two-tailed.

**Bulk RNA-sequencing extraction, alignment, and quantification**
RNA was extracted from adrenal tissues homogenized in TRIzol (15596026, Thermo Scientific) then processed using an RNeasy kit (74004, Qiagen). Total RNA libraries were prepared using the NEBNext® Poly(A) mRNA Library Prep Kit (E7770; New England Biolabs) to isolate polyadenylated transcripts. RNA integrity numbers ranged from 7.5–9.7. Libraries were sequenced using paired-end 150-bp reads (2×150 cycles) on the G4 sequencing platform (Singular Genomics, San Diego, CA, USA). High-throughput sequencing yielded a mean read depth of $5.04 \times 10^6$ reads per sample (median: $4.70 \times 10^6$; range: $2.95 \times 10^6$–$10.73 \times 10^6$). Sequence reads were aligned to the *Mus musculus* genome (mm10), achieving a high degree of mapping consistency with a mean genomic mapping rate of 96.34% (range: 95.86% – 97.52%). Reads were assigned to genes using ENSEMBL gene annotations. Following alignment, reads were assigned to genomic features; an average of 74.27% of total reads per sample (range: 71.85% – 76.28%) were successfully assigned to specific gene features for downstream differential expression analysis. Gene expression quantification was performed on the resulting BAM files using the *featureCounts* function from the *Rsubread* package. Quantification was conducted in paired-end mode with multi-threading enabled. The resulting raw count matrix was exported for downstream analysis.

**Single-cell RNA-sequencing reference processing**
To facilitate cell-type deconvolution, a single-cell RNA-seq (scRNA-seq) reference dataset was compiled from publicly available adrenal gland data (GEO Accession: GSM4914026 for Control, GSM4914027 for Stressed). The processing, clustering, and annotation of the single-cell reference data followed the methodology described in the original source publication. Raw sparse matrices were loaded into R (4.3.3) and processed using the *Seurat* package. Control and Treated datasets were merged, and cells were filtered based on quality control metrics (200–6,000 genes detected; <20% mitochondrial content). The combined dataset underwent normalization, variance stabilization, and scaling. To correct for batch effects between conditions, the ComBat algorithm from the *sva* package was applied. Dimensionality reduction was performed using PCA, followed by unsupervised Louvain clustering (resolution 0.5). Clusters were annotated into major cell types (e.g., Zona glomerulosa, Zona fasciculata, Macrophages) consistent with the marker genes and definitions established in the original study.

**Bulk RNA-seq deconvolution**
Cell type proportions in the bulk RNA-seq samples were estimated using the MuSiC (Multi-subject Single Cell deconvolution) method (*78*). The processed scRNA-seq object was converted to a *SingleCellExperiment* format to serve as the reference. ENSEMBL IDs in the bulk dataset were mapped to gene symbols using the org.Mm.eg.db annotation database to ensure compatibility with the single-cell reference. The *music_prop* function was used to estimate the weighted proportion of each cell type in the bulk samples.

**Differential expression analysis**

Differential expression analysis was performed using DESeq2 on the combined male and female bulk RNA-seq dataset (*79*). To account for sex and adrenal cell-type composition, the DESeq2 design formula included sex, the estimated zona fasciculata cell proportion from MuSiC deconvolution, and treatment condition: ~ sex + zf_prop + treatment. Differential expression was tested for fentanyl-treated adrenal glands relative to saline-treated controls, with positive log2 fold-change values indicating higher expression in fentanyl-treated samples. Log fold-change shrinkage was performed using the apeglm method. Differentially expressed genes were identified using a Benjamini–Hochberg adjusted p-value cutoff of $< 0.05$ and a log2 fold change threshold of $|2|$.

**Targeted HPA-Axis Gene Expression Analysis**

To evaluate treatment effects on hypothalamic-pituitary-adrenal (HPA) axis function, we performed a focused analysis on 8 HPA axis genes: *Mc2r, Mrap, Nr3c1, Fkbp5, Pomc, Crhr1, Crhr2,* and *Crhbp*. Gene expression values were extracted from the featureCounts-derived count matrix and normalized to $\log_2(\text{CPM} + 1)$. We performed a two-way ANOVA with gene and treatment group (fentanyl-treated vs. saline-treated controls) as fixed factors. This approach partitions variance into gene-specific effects (expected to dominate) and treatment effects while testing for gene-by-treatment interactions. A significance threshold of $\alpha = 0.05$ was applied. The analysis included all individual expression measurements across both sexes ($n = 192$ observations: 14 untreated samples × 8 genes + 10 treated samples × 8 genes).

**Functional enrichment analysis**

Gene Ontology (GO) enrichment analysis was performed on genes upregulated or downregulated in fentanyl-treated adrenal glands using the *clusterProfiler* package (*80*). Enrichment was restricted to Gene Ontology Biological Process (BP) terms and corrected for multiple hypothesis testing using the Benjamini–Hochberg method, with $q < 0.05$ considered significant. Upregulated and downregulated BP terms are shown in the main figures. Bulk RNA-seq deconvolution results and gene-concept network plots for upregulated and downregulated gene sets are shown in the supplemental figures.

**Solution preparation**

Drug solutions were prepared fresh or from stock as appropriate on the day of administration. For injection experiments, crinecerfont and fentanyl were first dissolved in dimethyl sulfoxide (DMSO) to generate concentrated stock solutions, which were subsequently diluted to the final working concentration in sterile 0.9% saline prior to injection. ACTH, naloxone hydrochloride, and naloxone methiodide were prepared directly in sterile saline at their respective working concentrations. Saline control animals received sterile 0.9% sodium chloride vehicle. Specific doses and administration routes are detailed in Table S2. For vapor-based self-administration assays, fentanyl was dissolved at 1 mg/mL in a vehicle consisting of 80% vegetable glycerin (Essential Depot) and 20% propylene glycol (02151957, MP Biomedicals) as previously described (*58*, *60*). Care was taken to ensure that final injection solutions were thoroughly mixed and visually free of particulates before administration.

**Stereotaxic craniotomy, viral injection, and implantation**

All surgical procedures were performed under aseptic conditions in accordance with institutional guidelines. Crh-IRES-Cre or C57BL/6 mice aged 6-10 weeks were anesthetized with isoflurane (0.5–2.5% in $O_2$) and secured in a stereotaxic frame (David Kopf Instruments) on a temperature-controlled heating pad. Ophthalmic ointment was applied to prevent corneal drying. Following induction of anesthesia, the scalp was disinfected and a midline incision was made to expose the skull. Connective tissue was cleared, and the skull was leveled by aligning bregma and lambda according to a mouse brain atlas (*81*). Stereotaxic coordinates were defined relative to bregma. A small craniotomy was drilled over the target with coordinates as follows: nucleus accumbens (NAc) ML -1.4, AP+1.2, DV -4.5; paraventricular nucleus of hypothalamus (PVN) ML -0.25, AP -0.8, DV -4.5; and ventral tegmental area (VTA) ML -0.45, AP -3.0, DV -4.5. A microsyringe (NanoFil syringe coupled to a UMP-3 pump, World Precision Instruments) was used to deliver a viral vector at a titer of $\geq 1\times10^{13}$ vg/mL. Half of the volume was injected 0.1 mm below the DV coordinate and the other half at exactly the DV coordinate at a rate of ~150–300 nL·min$^{-1}$. Following each injection, the needle or implant was left in place for ~10 minutes to allow for diffusion and minimize backflow.

For multisite fiber recordings, virus was injected into both the NAc (400 nl) and PVN (400 nl) and 2 weeks later, the device was implanted into the NAc, PVN and VTA. For DREADD experiments, virus was injected into both the NAc (400 nl) and PVN (400 nl), and a 400 µm silica fiber (FT400UMT) was implanted into the NAc. For PVN recording experiments, 400 nl of virus was injected into the PVN, and either a 400 µm silica fiber (FT400UMT) or multifunctional fiber was implanted into the PVN. Surgical implantations for electrophysiology experiments included a stainless-steel ground screw as well as a second reference screw secured to the skull over the cerebellum. The implant was affixed using adhesive (C&B Metabond, Parkell) followed by dental cement (Jet-Set 4, Lang Dental). The incision was closed around the implant base, and animals received post-operative care including monitoring and recovery on a heated surface. Mice were singly housed post-surgery under a 12 h light/dark cycle with ad libitum access to food and water. These surgical procedures follow established protocols for combined viral transduction and optical fiber implantation optimized for fiber photometry recordings in freely-behaving mice (*82*, *83*).

**Local field potential data acquisition and spectral analysis**

Local field potential (LFP) recordings were obtained from the 20 µm tungsten wire electrodes (Goodfellow) of the multifunctional neurophysiology fiber devices using Ripple Neuro acquisition system (Ripple, Salt Lake City, UT) and stored as NS2 files. Data were imported into MATLAB (MathWorks, 2022b) using the Neural Processing MATLAB Kit (NPMK), with sampling frequencies extracted from file metadata for each recording (*82*). Analyses were restricted to a predefined subset of electrodes per device and condition to ensure consistency across sessions.

Raw signals were oriented into channel × time format. Signals were bandpass filtered between 1–200 Hz using a fourth-order Butterworth filter (*zero-phase, filtfilt*), and narrowband noise was removed using sequential notch filters centered at 28, 60, 86, and 180 Hz. Time–frequency representations were computed using short-time Fourier transforms (STFT; 2 s window, 75% overlap, NFFT = 2×window length), and spectral power was calculated as the squared magnitude of the complex spectrogram. Analyses were restricted to frequencies between 0–200 Hz. For aggregate spectrograms, linear power was averaged across channels and devices within each condition and converted to decibel units. A global color scale (5th–95th percentile across all conditions) was applied to enable direct comparison. Residual narrowband noise was visually attenuated by interpolation across notch-filtered frequencies for display purposes only. Spectral analyses were performed using Chronux (*84*, *85*). For acute within-mouse analyses shown in Fig. 1, gamma power was subdivided into low gamma [30–80 Hz) and high gamma [80–200 Hz). For delayed between-group PVN LFP analyses shown in Fig. 3, frequency bands were defined as delta [1–4 Hz), theta [4–8 Hz), alpha [8–13 Hz), beta [13–30 Hz), and gamma [30–200 Hz).

**Band Power Quantification and Statistical Analysis**

Spectrograms were computed for each electrode and averaged across electrodes and time to obtain a single power spectrum per recording. For band-resolved analyses, frequency bands were defined as specified above for each experiment. Band power was calculated as the mean power spectral density within each band. For comparisons across independent groups (control vs treatment; Fig. S2), band power was expressed in decibel units ($10 \cdot \log_{10}$ power/Hz) and analyzed using a mixed-effects framework with Band as a within-subject factor and Group as a between-subject factor (*86*). Device-level values were computed by averaging across electrodes within each recording. For within-recording comparisons across behavioral epochs (baseline vs Straub; Fig. 1), band power was normalized within each device by subtracting baseline values, yielding baseline-normalized changes in power (ΔdB). These values were used for both visualization and statistical analysis. Statistical comparisons for within-recording analyses were performed using a repeated-measures ANOVA with within-subject factors of Band and Epoch (baseline vs Straub), implemented using MATLAB's *fitrm* and *ranova* functions. Greenhouse–Geisser correction was applied to account for violations of sphericity (*87*, *88*). Significant main effects and interactions were followed by post hoc paired t-tests on baseline-normalized changes (ΔdB) within each frequency band. Resulting p-values were corrected for multiple comparisons using the false discovery rate (FDR) method (*89*).

**AAV packaging**

Adeno-associated virus serotype 9 (AAV9) carrying Cre-dependent hM4D(Gi) fused to fluorescent protein mCherry (AAV9-hSyn::DIO-hM4D(Gi)-mCherry, titer $1.7 \times 10^{13}$ vg/mL) was produced following polyethyleneimine (PEI)-mediated triple transfection protocol (*90*). Briefly, plasmids pAAV2/9n (Addgene 112865), pAAV- hSyn::DIO-hM4D(Gi)-mCherry (Addgene 44362), and pHelper (CELL BIOLABS, Inc.) were used for PEI-mediated triple transfection of HEK293T cells. The culture medium was collected at 72 and 120 h, and the cells were harvested at 120 h post-transfection, followed by purification by ultracentrifugation. The AAVs were collected in Dulbecco's phosphate-buffered saline (DPBS; D8662, Gibco) with 0.001% Pluronic F-68 (Gibco). AAV titers were determined using a Takara Bio AAV real-time PCR titration kit (#6233). Viral preparations were stored in aliquots at −80°C until use.

**Fiber photometry acquisition**

Animals were anesthetized with 1% isoflurane, placed on a heated surface, and ophthalmic ointment (Puralube) was applied to the eyes. Recordings were performed using a Neurophotometrics FP3002 system, which enables simultaneous dual-wavelength acquisition from multiple brain regions in freely moving animals. LEDs were calibrated to deliver 100 μW of optical power from the fiber tip, and recordings were acquired at 130 Hz, with alternating 470-

nm (experimental channel) and 415-nm (reference/isosbestic channel) excitation (*91*). The optical fiber was pre-bleached with high-intensity illumination for 1 hour before each photometry recording session to eliminate residual autofluorescence from the patch cord. For naloxone pretreatment experiments, mice received naloxone hydrochloride (BP548, Sigma-Aldrich; 5 mg·kg$^{-1}$, s.c.) 400 s before fentanyl administration. Control mice received an equivalent volume injection of saline before fentanyl.

**Chemogenetic PVN$^{CRH}$ inhibition and imaging**

For chemogenetic inhibition of PVN$^{CRH}$ neurons during fentanyl administration, Crh-IRES-Cre mice (n = 4; 2 males, 2 females) and age-matched C57BL/6 control mice (n = 4; 2 males, 2 females) received 400 nL injections of AAV9 carrying Cre-dependent inhibitory DREADD into the PVN (1.7 × $10^{13}$ vg/mL; pAAV9-hSyn-DIO-hM4D(Gi)-mCherry; 44362, Addgene) and 400 nL of AAV9 carrying dLight1.3b into the ipsilateral NAc (≥ 1 × $10^{13}$ vg/mL; pAAV9-syn-dLight1.3b; 135762, Addgene) (*92–94*). A 400-µm silica optical fiber (FT400UMT, Thorlabs) was implanted into the NAc for dopamine photometry recordings using the Neurophotometrics FP3002 system. During recordings, mice were maintained under light anesthesia with 1% isoflurane and received clozapine N-oxide (CNO; 4936, Tocris; 10 mg·kg$^{-1}$, s.c.) 10 min before fentanyl administration (0.4 mg·kg$^{-1}$, s.c.; 1269902, Sigma-Aldrich). Calculations for %$\Delta F/F_0$ were taken from 200 s epochs approximately 4.5 min following fentanyl injection.

**Fiber photometry analysis**

Fiber photometry data were analyzed from preprocessed fluorescence recordings imported into Python (3.12) (*NumPy, Pandas, SciPy*). A second-order zero-phase Butterworth low-pass filter (cutoff frequency = 10 Hz) was applied to the raw 470-nm signal to reduce high-frequency noise and electrical artifacts, followed by a second-order zero-phase Butterworth high-pass filter to correct for photobleaching drift and low-frequency baseline fluctuation (*95*). Zero-phase forward-backward filtering (*filtfilt*) was used throughout to eliminate temporal phase distortion and preserve the timing of dopamine transients (*96*). The 415-nm isosbestic control channel was acquired simultaneously to validate the absence of motion artifacts during recordings; given that all experiments were conducted under isoflurane anesthesia with minimal subject movement, the 415-nm channel served primarily as a motion-artifact control validation rather than active artifact correction. Data preprocessing initially generated z-scored fluorescence values; these z-scores were subsequently back-converted to %$\Delta F/F_0$ according to the transformation: %$\Delta F/F_0$ = (z-score × std) + mean, where the mean and standard deviation were calculated from the preprocessed filtered signal. All quantitative statistical analysis was performed on the resulting %$\Delta F/F_0$ values. Following normalization, traces were smoothed using a moving-average filter (window = 65 samples, ~0.5 s at 130 Hz) and baseline-centered then shifted uniformly using a pre-injection epoch to normalize signals across subjects and avoid negative baseline values (*97*). Excessive smoothing was avoided to ensure that peak amplitude, rise time, and decay kinetics of dopamine transients were appropriately preserved.

Predefined time windows (epochs) aligned to experimental events were extracted for each animal. Within each epoch, the mean %$\Delta F/F_0$ and standard error of the mean (SEM) were calculated from the preprocessed, smoothed signal. For pairwise group comparisons (e.g., control vs. naloxone-treated; C57BL/6 vs. CRH-Cre), a two-way ANOVA was performed. For multi-epoch comparisons within a single group (e.g., baseline vs. saline vs. post-fentanyl), one-way ANOVA was performed followed by Dunnett's post-hoc test to compare each treatment condition against the baseline control. Dunnett's test was selected to control family-wise Type I error rates while maintaining statistical power for control-vs-treatment comparisons. All statistical analyses were conducted in GraphPad Prism.

**Fluorescence immunostaining**
Mice were euthanized by intraperitoneal administration of Fatal-Plus (100 mg·kg$^{-1}$), followed by transcardial perfusion with ice-cold phosphate-buffered saline (PBS; 46-013-CM, Corning) diluted to 1X and 4% paraformaldehyde (PFA; 15714-S, Electron Microscopy Sciences). Brains were extracted, post-fixed in 4% PFA for 24 h at 4°C, and transferred to PBS. Coronal sections (40 µm) were prepared using a vibratome (Leica VT1000S). GCaMP6s and hM4D(Gi)-mCherry fluorescence from prior virus injections were imaged directly without antibody amplification, as the endogenous signal from viral vector expression was sufficient for visualization. For dLight1.3b and c-Fos visualization, sections were permeabilized in 0.3% Triton X-100 (X100, Millipore Sigma) in PBS and blocked in 0.3% Triton X-100 with 3% bovine serum albumin (A215, Millipore Sigma). Sections were incubated with rabbit anti-GFP primary antibody (A-11122, Thermo Scientific; 1:200) for dLight1.3b or anti-c-Fos Rabbit Monoclonal Antibody [9F6] (Cell Signaling Technology, 2250S; 1:200) for 48 h at 4°C. After 3 PBS washes, sections were incubated with Alexa Fluor 488-conjugated secondary antibody specific to rabbit IgG (ab150077; 1:1,000) for 1 h. All sections were counterstained with DAPI (1:10,000; 62248, Thermo Scientific), washed, and mounted onto glass slides using Fluoromount-G (0100-01, Southern Biotech) with No. 1.5 coverslips (*98*).

**Confocal imaging**
Images were acquired using a Leica Stellaris 5 confocal microscope equipped with a white light laser (WLL, 485–790 nm excitation) and Power HyD photon-counting detectors. This configuration enables the freedom to select multiple simultaneous excitation wavelengths and detection windows optimized for the spectral profiles of all fluorophores in the sample, up to 8 independent laser lines. Images were acquired with a HC PL APO 10×/0.40 CS2 objective (Leica part #11506424). Regions of interest were selected based on implant locations and viral expression patterns.

**cFOS+ neuronal quantification**
For each image, triangular regions of interest (ROI) encompassing the PVN were manually defined based on anatomical landmarks using Python scripts (scikit-image). DAPI ($\lambda$=405 nm) and cFOS immunofluorescence ($\lambda$=488 nm) channels were cropped to their respective ROI regions and saved as separate grayscale TIFF files. Image analysis was conducted manually in FIJI (ImageJ) (*99*). Nuclei were identified in the DAPI channel using automatic thresholding (Otsu) followed by particle analysis (Analyze Particles). cFOS+ nuclei were identified by visual inspection of overlapping signal in the cFOS channel within detected nuclear regions. The percentage of cFOS+ nuclei was calculated as (cFOS+ nuclei / total nuclei) × 100.

**Jugular vein catheterization**
Jugular vein catheterization was performed under aseptic conditions. Briefly, mice were anesthetized with isoflurane (3% induction, ~2% maintenance) and placed on a heated surgical surface. Dorsal and ventral incision sites were shaved, chemically depilated, and sterilized using alternating applications of betadine and ethanol. A midline dorsal incision and a right-sided

ventral cervical incision were made to expose the external jugular vein. A subcutaneous tunnel was created between incision sites, and a polyurethane catheter (C20PU-MJV1301, Instech) was routed from the dorsal access site to the ventral incision. The jugular vein was isolated, and two ligatures were placed rostral and caudal to the insertion site. A small venotomy was made, and the catheter was inserted into the vein and advanced to a premeasured depth. The catheter was secured using suture ties, and patency was confirmed by aspiration of blood followed by saline flushing. The catheter was exteriorized and connected to a vascular access port (VABM1B/25) secured at the dorsal incision site and protected with a magnetic cap (VABM1C). Animals were allowed to recover on a heated pad and monitored until fully ambulatory. Catheters were subsequently accessed using sterile injectors (PNP3M) and appropriate handling tools (VABMG) for maintenance and sampling (*100*).

**Acute ACTH, β-endorphin, and corticosterone measurements by submandibular puncture**
Blood was collected for the following experiments by submandibular puncture using Goldenrod animal lancets (5 mm; Braintree Scientific). Blood was collected into lithium heparin tubes (MV-100 201282-BX, Braintree Scientific), and plasma samples were stored at −80°C until analysis (*101*).

For acute ACTH and β-endorphin measurements, blood was collected 20–30 min after saline or fentanyl injection. Circulating ACTH was measured using an ACTH ELISA kit (ab263880, Abcam), and β-endorphin concentrations were measured using a β-endorphin ELISA kit (EEL087, Fisher Scientific), according to the manufacturers' instructions.

For ACTH stimulation testing, 6-week-old C57BL/6 mice were treated once daily for 4 days with fentanyl (0.4 $mg \cdot kg^{-1}$, i.p.) or saline. On day 5, mice received ACTH (A0298, Sigma-Aldrich; 1 $mg \cdot kg^{-1}$, i.p.), and blood was collected after ACTH administration to assess stimulated corticosterone release (*102*). Sample collection across the cohort occurred over an 80-min period, and each mouse was plotted according to its blood collection time relative to ACTH administration. Plasma corticosterone concentrations were measured using a corticosterone ELISA kit (ab108821, Abcam) according to the manufacturer's instructions. Corticosterone concentrations were plotted as a function of elapsed time after ACTH injection, and simple linear regression with 95% confidence intervals was used to visualize stimulated corticosterone dynamics within each treatment group.

For assessing circulating corticosterone following adrenal gland hypertrophy, mice were treated once daily for four days with ACTH (A0298 Sigma-Aldrich; 0.45 $mg \cdot kg^{-1}$, i.p.) or saline. Plasma corticosterone concentrations were measured after the pretreatment period using a separate corticosterone ELISA kit (ab323743, Abcam) according to the manufacturer's instructions.

For acute ACTH and β-endorphin measurements, between-group comparisons (saline vs. fentanyl) were performed using Welch's t-test. For ACTH stimulation testing, corticosterone responses were visualized using simple linear regression with 95% confidence intervals, and Welch's t-test was used to compare the slopes and intercepts between treatment groups. For circulating corticosterone following adrenal gland hypertrophy, Welch's t-test was used to compare ACTH and saline-treated groups. All statistical analyses were conducted in GraphPad Prism (11.1.0).

**Longitudinal ACTH measurements during acute fentanyl exposure**
For longitudinal ACTH measurements during acute fentanyl exposure, jugular-catheterized male mice were sampled at baseline and at 20, 40, 60, 80, and 100 min after saline or fentanyl

administration with samples taken from the same mice between 14:00-15:00 on sequential days. Circulating adrenocorticotropic hormone (ACTH) levels were measured using a commercially available ELISA kit (ab263880, Abcam) according to the manufacturer's instructions. ACTH values from longitudinal sampling experiments were normalized to each subject's baseline and expressed as percent baseline. Data were analyzed in GraphPad Prism (11.1.0) using two-way repeated measures ANOVA with treatment (saline vs. fentanyl) as the between-subjects factor and time point (0, 20, 40, 60, 80, 100 min) as the within-subjects factor. Data are presented as mean ± SEM.

**HPA axis gland mass model**

HPA axis dynamics were simulated in Python (3.11.15) using a reduced gland-mass model adapted from Milo et al. (*49*, *50*) The model links fast hormone dynamics to slower changes in adrenal functional mass. Stress input drives corticotropin-releasing hormone (CRH) activity, CRH stimulates adrenocorticotropic hormone (ACTH) release, and ACTH promotes corticosterone production in proportion to adrenal functional mass. Corticosterone provides negative feedback onto upstream CRH and ACTH signaling. To focus on adrenal remodeling, the pituitary functional-mass variable from the original model was removed, while ACTH was retained as a dynamic hormone variable.

The model consisted of coupled ordinary differential equations describing CRH, ACTH, corticosterone, and adrenal functional mass. Adrenal functional mass changed according to ACTH-dependent adrenal cell-mass production and adrenal cell-mass removal. Fentanyl was incorporated as a time-dependent perturbation to hypothalamic input, modeled as an initial transient increase in CRH drive followed by sustained inhibition. Exogenous ACTH administration was simulated by adding an external ACTH input during the pretreatment period. Simulations were initialized at baseline steady state before introduction of experimental perturbations, and model outputs were expressed relative to baseline. Numerical integration of the ODE system was performed in Python using SciPy solvers.

**Fentanyl vapor chamber design and fabrication**

The system was adapted from previously validated vapor-based self-administration paradigms and further engineered in consultation with the MIT Environmental Health and Safety (EHS) to ensure safe handling and containment of aerosolized opioids (*58*–*60*). The operant chamber was designed to be airtight, minimizing vapor leakage and ensuring precise control over drug delivery and clearance. The system consisted of an operant chamber equipped with active and inactive nose-poke ports (ENV-375A-NPP, Med Associates), a grid floor capable of delivering foot shock (ENV-307W-GFW, Med Associates), and a resistive heating coil (0.2 Ω, 70–80 W; Geekvape) for drug vaporization. Behavioral events, including nose-pokes and stimulus contingencies, were controlled and recorded using Med Associates hardware and software. Upon completion of the required number of active nose-pokes, the coil was activated to vaporize the fentanyl solution, which was delivered into the chamber via a tightly regulated airflow system driven by a vacuum pump. Activation duration (5 s) was optimized to ensure reproducible dosing and rapid clearance, with vapor levels returning to baseline within approximately 30 seconds between deliveries. To further ensure containment, exhaust air was passed through an inline HEPA filter and routed through a biosafety cabinet prior to entering the facility ventilation system.

**Fentanyl Self-Administration Assays**

Mice were pretreated with adrenocorticotropic hormone (ACTH; 0.45 $mg \cdot kg^{-1}$, i.p.) or saline once daily for 4 consecutive days and compared to saline-treated littermate controls. Operant fentanyl self-administration was conducted in custom vapor-delivery chambers equipped with active and inactive nose-poke ports. Behavioral sessions were controlled using Med Associates hardware and custom software implementing a fixed-ratio (FR) schedule of reinforcement (FR1 unless otherwise specified) (*103*). Each session lasted 30 minutes, with a maximum of 40 reinforcers permitted. Active nose-pokes triggered fentanyl vapor delivery, followed by a timeout period (~45 s) during which additional responses were recorded but did not result in drug delivery. Inactive nose-pokes were recorded but had no programmed consequence. Behavioral metrics, including total active and inactive nose-pokes, timeout responses, and total reinforcers, were recorded in real time. The behavioral paradigm consisted of three sequential phases. During acquisition (days 0–4), mice were trained under an FR1 schedule to discriminate the active nose-poke, with preference defined as >60% active responses. During the effort escalation phase (days 5–7), the response requirement was increased to a fixed ratio of three (FR3). During the punishment phase (days 8–10), active nose-pokes and drug delivery were paired with a foot shock of increasing intensity (0.1, 0.5, 0.7 mA; 0.5s duration) across days. Active nosepokes from Days 5–10 (FR3 and FS phases) were analyzed using two-way ANOVA with Group (Saline vs. ACTH) and Day as fixed factors, with Type II sums of squares. For phase-level comparisons, active nosepokes and related metrics were collapsed across days within each phase (FR3: Days 5–7; FS: Days 8–10) for each mouse, and group means were compared using independent samples t-tests with Welch's correction. Cohen's d was calculated for all pairwise comparisons to quantify effect magnitude. Partial eta squared ($\eta^2$) was reported for ANOVA effects. During acquisition (FR1, Days 0–4), percentage active nosepokes were compared day-by-day using t-tests with Benjamini–Hochberg FDR correction (*89*, *104*, *105*). All tests were two-tailed. Analyses were performed in Python (3.12) using *pandas, SciPy, and statsmodels*.

**Multisite imaging fiber fabrication**

Multi-site imaging fiber devices were designed to interrogate specific brain regions of interest. A high-resolution 3D printer (Stratasys J5 Medijet Polyjet 3D printer) was used to print an enclosure with holes positioned to access brain regions of interest: NAc: ML -1.4, AP+1.2, DV -4.5; PVN: ML -0.25, AP -0.8, DV -4.5; VTA: ML -0.45, AP -3.0, DV -4.5. Holes centered at the NAc, PVN, and VTA positions were printed at a diameter of 0.55 mm, 0.55 mm, and 0.3 mm, respectively, to accommodate each waveguide. To image the PVN and VTA with spatial resolution, a multi-core polymer optical fiber (Asahi Kasei) was used, containing 7400 independent 4 µm waveguides within a 500 µm footprint. To detect neurotransmitter release in the NAc, a 250 µm polymer optical fiber (Mitsubishi) was used. Each fiber was threaded through its respective hole in the 3D-printed enclosure and a custom 3D-printed ferrule and secured in place with epoxy (Norland Optical Adhesive) cured under ultraviolet light. Individual fibers were trimmed to appropriate lengths and polished manually at both ends. The spatial mapping of each fiber between its location in the ferrule at the top (output side) of the device and its location at the bottom (implant side) of the device was confirmed by capturing images of the output side while translating the position of a fluorescent target under each of the three individual fiber facets on the implant side. Following implantation, the external fiber ports were protected using dust caps to minimize the accumulation of debris.

**Multisite imaging and data analysis**
A custom epifluorescence microscope system was used to simultaneously record imaging data through each of the three fiber facets. A fiber-coupled LED (Thorlabs, M470F3) was used to provide excitation at 470nm following an excitation filter (470/22nm, Semrock BrightLine FF01-470) through a 10X objective lens (Olympus, UPLXAPO). A single-edge dichroic beamsplitter (495nm, Semrock BrightLine FF495-Di03) was used to route emitted fluorescence through emission filters (488nm long-pass, Semrock BrightLine LF488; 525/50nm, Semrock BrightLine FF03-525) to a high-speed camera (FLIR, Blackfly BFS-U3-200S6M-C) operated at 10 Hz (0.1 s exposure time) controlled by custom software for real-time visualization and data acquisition (Python). A tube lens of focal length 125mm (Thorlabs) was used for an effective magnification of 7.6X over a 900× 900 $\mu m^2$ field of view. Although the optical power incident on the device used was 24.0 mW, due to the distribution of the focused beam over an area encompassing all three fibers of the multi-site imaging device, the optical power output from all three fibers was measured to be 1.51 mW ± 0.03 mW (mean ± st.dev, n = 3 devices), which is consistent with the low reported loss of the polymer waveguides (0.0035 dB/cm for a single-core fiber, 0.018 dB/cm for multi-core fiber).
For multisite PVN, VTA, and NAc recordings, fluorescence signals were extracted from each fiber facet corresponding to each implanted region. For each region, the bulk fluorescence signal was averaged over the fiber facet and normalized to a background region to remove fluctuations in input light intensity. Signals were fit with a double exponential curve to correct for photobleaching, and relative fluorescence change at each pixel was quantified as $\Delta F/F_0$ relative to the initial baseline period. Mean normalized fluorescence was calculated during 2-min saline and fentanyl-evoked Straub epochs for each region and compared within mice using paired t-tests.

**Multifunctional neurophysiology fiber fabrication**
The twelve-electrode fiber with an optical waveguide was fabricated through a two-step thermal drawing process (*106*). First, a three-electrode fiber with dimensions of 50 × 200 $\mu m^2$ was created from a larger-scale preform consisting of two machined polycarbonate (PC, McMaster Carr) slabs, one with 1.575 mm channels. Teflon rods (McMaster Carr) were placed into the channels to hold them open while hot pressing the two slabs together at 175 °C and 5 psi for 30 min. The final preform dimensions were 16.75 × 4.19 mm. Following hot pressing, the Teflon rods were pulled out and the preform was drawn in a three-zone furnace set to temperatures of 125, 260, and 80 °C while converging three spools of 20 µm tungsten wire (Goodfellow). The feed rate was maintained at 0.25 m/min and capstan speed was set to ~1.6 m/min, resulting in a draw down ratio of ~84.
Four spools of the three-electrode fiber were then converged into a secondary preform, leading to a total of 12 electrodes. The secondary preform was made from multiple PC slabs with 4 channels machined to 12.7 × 3.18 mm for convergence of the three-electrode fiber, and a 14.29 mm diameter channel for the waveguide. Teflon spacers were used to hold the channels open while hot pressing PC slabs together at 180 °C and 5 psi for 60 min. A step-index waveguide with a core of polymethyl methacrylate (PMMA, McMaster Carr, nPMMA = 1.50) and THVP (terpolymer of tetrafluoroethylene, hexafluoropropylene, and vinylidene fluoride, 3M, nTHVP = 1.35) cladding was fabricated by wrapping a 12.7 mm PMMA rod in a 200 µm THVP film to a total diameter of 14.25 mm, encasing it in Teflon tape, and heating it at 160 °C for 30 min. The

Teflon was then removed, and the waveguide was inserted into the polycarbonate preform and consolidated at 100 °C overnight. The fiber was drawn in a three-zone furnace with temperatures set to 130, 255, and 100 °C while converging 4x spools of the three-electrode fiber. A feed rate of 0.25 m/min and capstan speed of ~1 m/min was used to achieve a draw down ratio of ~64 and final fiber size of ≤ 400 × 400 $\mu m^2$.

**Printed circuit board (PCB) manufacturing**

A custom single-layer PCB was fabricated to enable compact connectorization of the fiber. The PCB was patterned using an LPKF Protolaser U4 by laser ablating 18 µm copper on a 1.5 mm FR4 board (MG Chemicals). Thru-holes were also cut to enable pinning of electrodes and pass-through of the waveguide ferrule.

**Device electrode impedance measurements**

Electrode impedance was measured at 1 kHz immediately prior to each experiment using the Ripple Grapevine neural interface system. Each channel was tested in situ to verify signal integrity and electrode functionality before data acquisition. Impedance values were recorded for all implanted contacts, and channels with abnormal impedance values indicative of open circuits, shorts, or poor contact were noted and excluded from subsequent analysis when appropriate. These measurements were used as a routine quality-control step to confirm stable electrode performance across recording sessions.

**Single unit recording and analysis**

Extracellular recordings were acquired using a Ripple Neuro system (Ripple, Salt Lake City, UT) at 30 kHz sampling rate from 8 microelectrodes, with signals converted from binary NS5 format (1 count = 0.25 µV). Continuous voltage traces were preprocessed with complementary filtering: a FIR notch filter (1000th order, 59–61 Hz stopband) removed 60 Hz line noise, followed by a 2nd-order Butterworth bandpass filter (300–6000 Hz passband) to isolate action potentials while attenuating drift and high-frequency noise. Putative spikes were detected on each channel using an amplitude threshold set at mean ± 5 standard deviations, applied separately to positive and negative deflections and subject to a 1 millisecond refractory period constraint. For each detected spike, a ±2 millisecond window around the threshold crossing was extracted as the waveform snippet. Each waveform was characterized by peak-to-peak amplitude (maximum voltage minus minimum voltage within the window, measured in µV) and trough-to-peak width (temporal distance from the negative trough to the subsequent positive peak, measured in milliseconds)  (*107*). Spikes were retained only if amplitude fell between 20–200 µV and width between 0.5–2 milliseconds, ensuring only plausible action potentials were analyzed. Retained waveforms for each electrode were subjected to principal component analysis (PCA) to reduce dimensionality, with the first 3 principal components typically explaining ≥80% of total variance. Spike clusters were identified using k-means clustering (up to 3 clusters per electrode) applied to principal component scores with k-means++ initialization and 10 random replicates, retaining units with ≥10 spikes per cluster. To eliminate potential multiunit contamination and cross-electrode spike detection, all spikes from all clusters on all electrodes were pooled and spike times were rounded to the nearest millisecond; for spike times occurring within 1 ms of each other across channels/clusters, only the spike with the largest amplitude was retained. Spike rasters were generated by assigning each sorted unit a unique row position (labeled by channel and cluster ID) and plotting each spike time as a dot at coordinates (spike_time, unit_row), providing a visualization of temporal firing patterns across all simultaneously recorded neurons. All analyses were performed in MATLAB using custom scripts, the NPMK toolkit, and the Chronux signal processing library (version 2.12), with results exported to comma-separated value files for statistical analysis (*108*). Spike counts for each neuron were tabulated in a firing rate export file containing metadata columns (Neuron_ID, Electrode, Cluster, Total_Spikes, Mean_FR_Hz) followed by time-binned firing rate estimates at 1-second intervals (T0.5_s, T1.5_s, etc., in Hz). The Total_Spikes column represents the cumulative spike count for each unit across the entire recording, calculated as the sum of all spike occurrences after redundancy elimination. Mean firing rate (Mean_FR_Hz) was derived by dividing total spikes by total recording duration (in seconds). Time-binned firing rates were calculated by integrating the Gaussian-smoothed instantaneous firing rate estimate within each 1-second window, allowing verification of total spike counts and temporal spike distribution patterns across the recording session.

**Supplementary Figures**

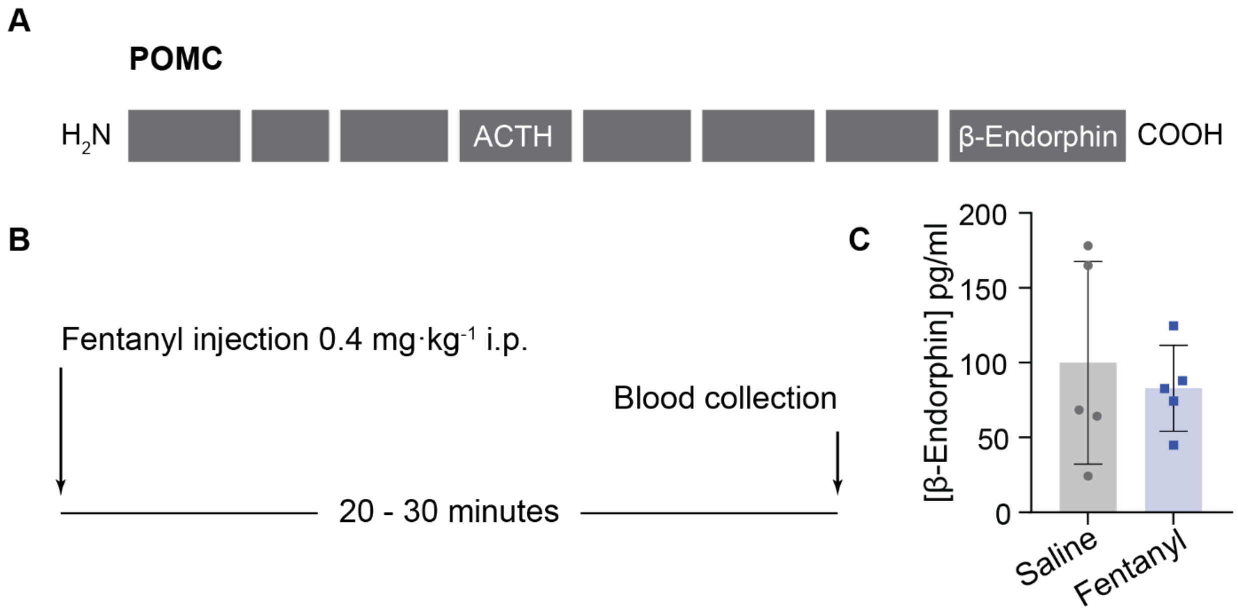


**Fig. S1 Fentanyl promotes desynchronization between HPA axis activation and endogenous opioid counter-regulation.** **(A)** β-endorphin and ACTH share the precursor peptide proopiomelanocortin (POMC). Cleavage sites indicated in white. **(B)** Plasma collected 20–30 min after saline or fentanyl injection. **(C)** Circulating β-endorphin concentrations (saline 99.96 ± 67.68; fentanyl 82.92 ± 28.66 pg/mL; Welch's $p = 0.6563$; $n = 5$ male mice per group). Bars and error bars represent mean ± standard deviation (SD).

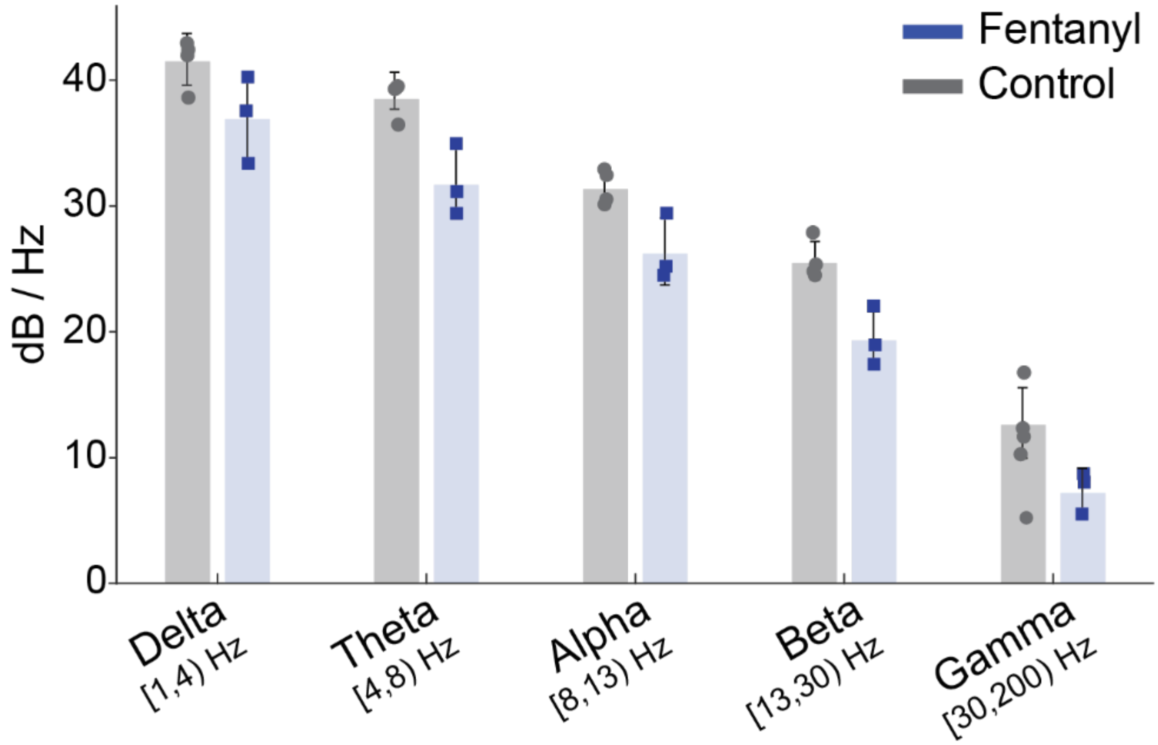


**Fig. S2 Fentanyl reduces PVN spectral power (dB/Hz) in the hours following administration.** Band-averaged spectral power recorded approximately 5 h after treatment was lower across all analyzed frequency bands in fentanyl-treated mice (n = 3; 2 males, 1 female) relative to control mice (n = 4; 2 males, 2 females). Delta (control: 41.47 ± 2.06 dB/Hz; fentanyl: 36.89 ± 3.46 dB/Hz), theta (38.49 ± 1.47 vs. 31.66 ± 2.84 dB/Hz), alpha (31.33 ± 1.38 vs. 26.20 ± 2.67 dB/Hz), beta (25.46 ± 1.54 vs. 19.30 ± 2.36 dB/Hz), and gamma power (12.58 ± 2.81 vs. 7.19 ± 1.79 dB/Hz). This broad reduction was reflected by a significant main effect of treatment group in a mixed ANOVA ($F(1,5) = 17.50$, $p = 0.00863$). Bars and error bars represent mean ± SD.

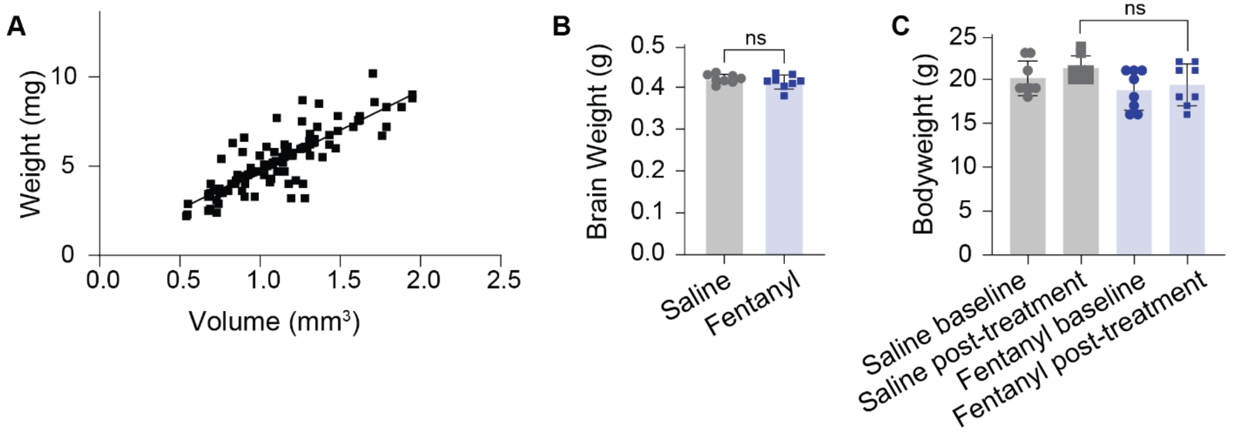


**Fig. S3 Validation of adrenal volume measurements and assessment of tissue and body weight after fentanyl exposure. (A)** Coefficient of determination between adrenal weight and volume $R^2 = 0.7225$ (n = 116 adrenal glands) **(B)** Mean and standard deviations of brain weights extracted from 7-week-old mice treated with four doses of fentanyl or saline (n = 8; Welch's p = 0.3122; 4 males, 4 females per group). **(C)** Mean and standard deviations of 7-week-old mouse body weights treated with four doses of fentanyl or saline (n = 8; Welch's p = 0.0864; 4 males, 4 females per group). **(B, C)** Bars and error bars represent mean ± SD.

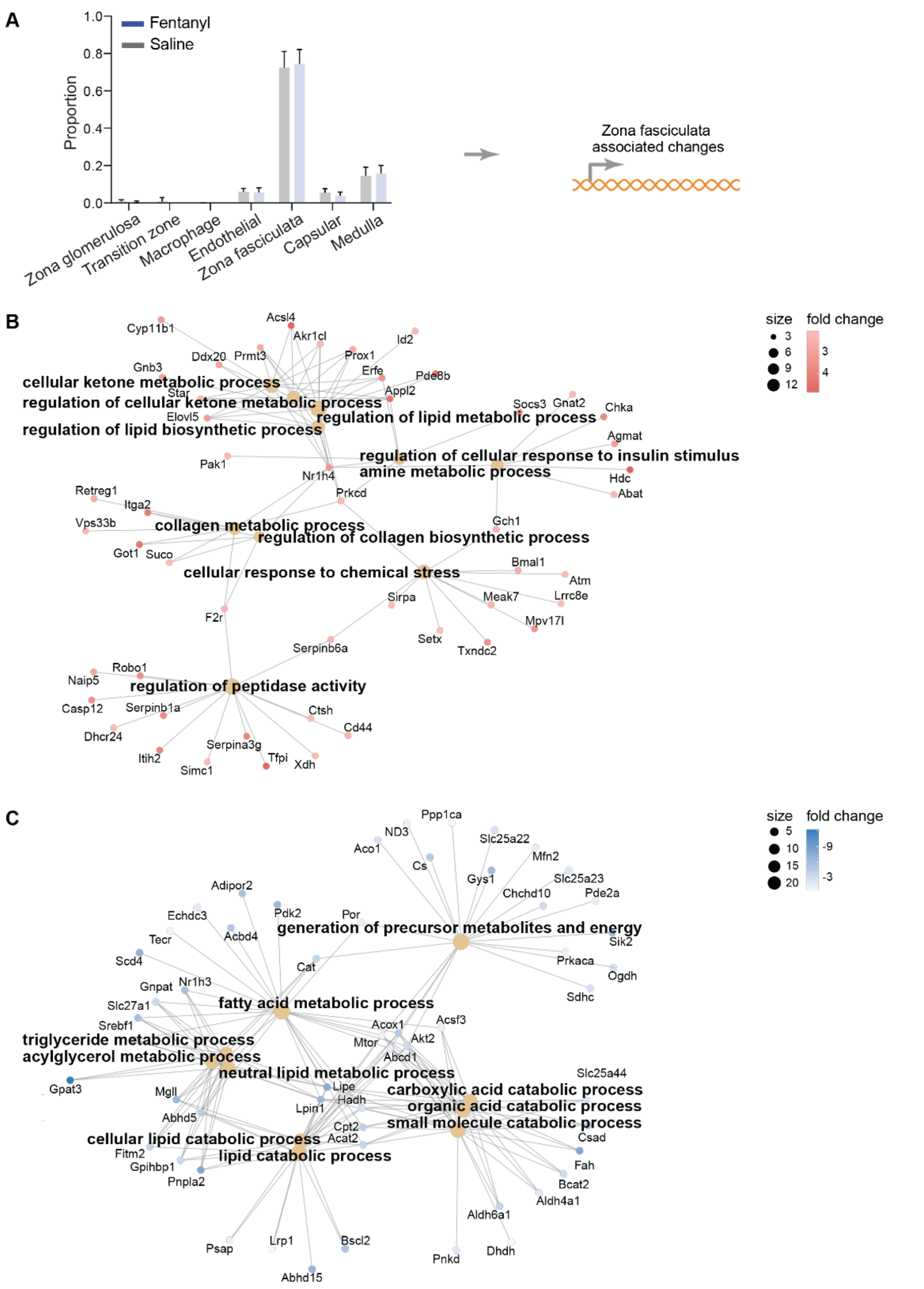


**Fig. S4 Fentanyl induces cellular stress and blunts normal metabolic processes within inferred zona fasciculata populations. (A)** Deconvolution estimates showing cell type proportion (fentanyl-treated (n = 10; 2 females, 8 males); saline-treated (n = 14; 8 females, 6 males). **(B, C)** Concept network (cnet) plots of predicted zona fasciculata-associated differentially expressed genes, depicting the relationships between significantly enriched biological pathways and their constituent genes. Node size reflects the number of genes associated with each enriched term, and gene nodes are colored by $\log_2$ fold change to indicate the direction and magnitude of differential expression in response to fentanyl.

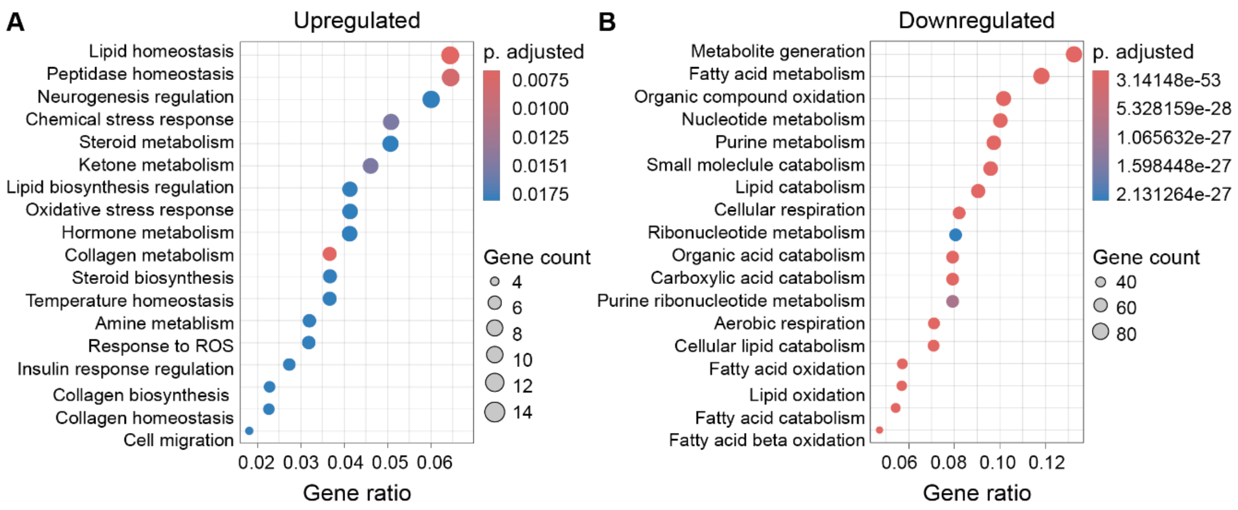


**Fig. S5 Gene Ontology Biological Process (GO BP) enrichment analysis of cell-type-specific differentially expressed genes resolved via bulk deconvolution.** Panels show processes enriched among upregulated **(A)** and downregulated **(B)** genes mapping to the zona fasciculata population in fentanyl-treated adrenal glands relative to saline-treated controls following Benjamini–Hochberg correction for multiple hypothesis testing ($q < 0.05$).

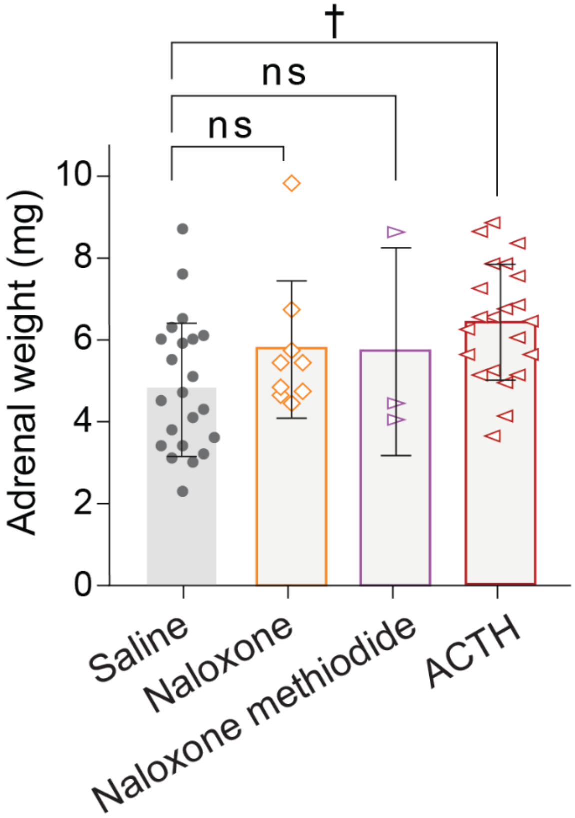


**Fig. S6 Adrenal weights in response to opioid antagonists and ACTH treatment.** Mean ± SD adrenal weights across treatment conditions. Saline control (n=12, mean = 4.88 ± 1.69 mg), naloxone (n = 5, mean = 5.77 ± 1.55 mg; p = 0.9986), naloxone methiodide (n=2, mean = 5.35 ± 1.48 mg; p = 0.7925), and ACTH (n=11, mean = 6.38 ± 1.4365 mg; p = 0.0833). Dunnett's test vs. saline. Bars and error bars represent mean ± SD.

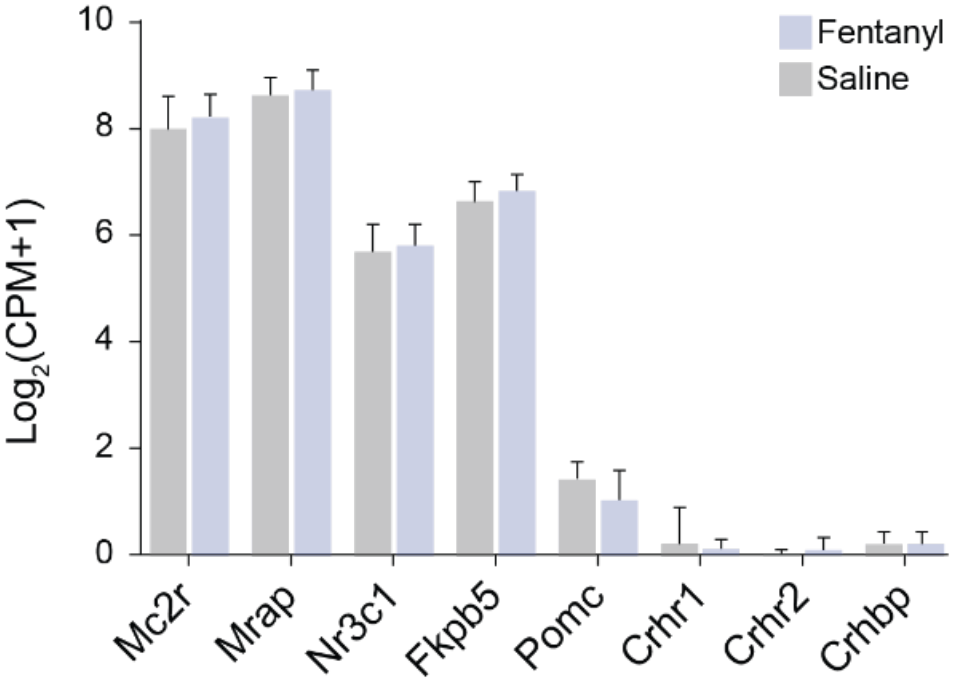


**Fig. S7 Stress-associated HPA axis gene expression in bulk adrenal tissue following fentanyl administration.** Mean + SD relative gene expression ($log_2$(CPM+1) for eight HPA axis genes in adrenal tissue from saline (n = 14; 8 females, 6 males) and fentanyl-treated (n = 10; 2 females, 8 males) mice. Genes include ACTH receptor components (Mc2r, Mrap), glucocorticoid receptor pathway (Nr3c1, Fkbp5), and CRH signaling (Pomc, Crhr1, Crhr2, Crhbp). Two-way ANOVA revealed a significant main effect of gene ($F[7,176] = 2008$, $p < 0.0001$) but no significant effect of fentanyl treatment ($F[1,176] = 0.276$, $p = 0.5999$). No significant gene-by-treatment interaction was observed ($F[7,176] = 1.446$, $p = 0.1897$). Bars and error bars represent mean + SD.

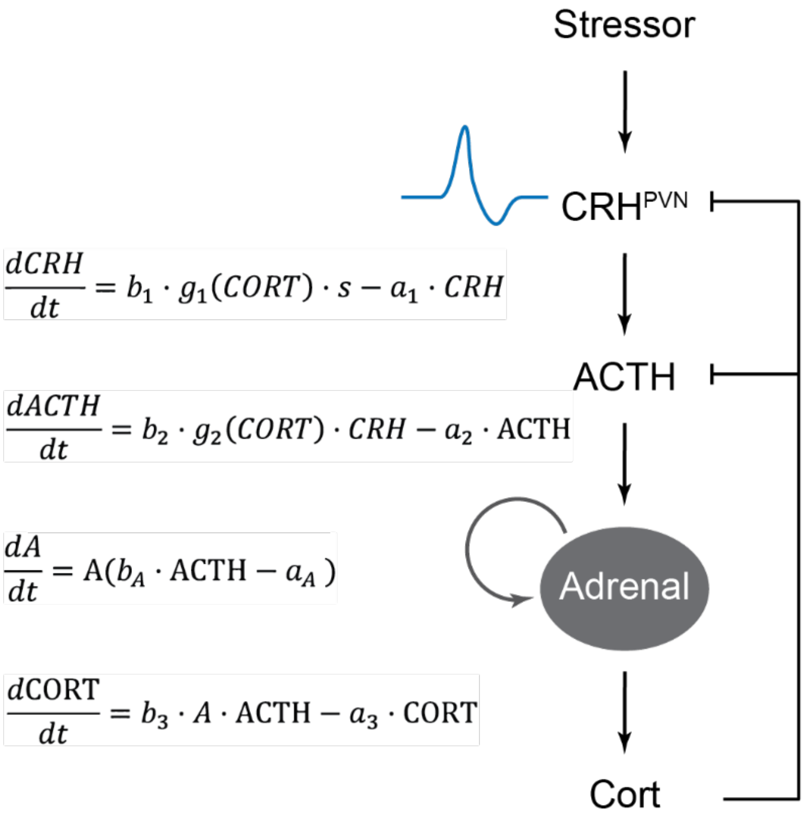


**Fig. S8 HPA axis model predictions reveal adrenal growth as a memory mechanism for prior stress exposure that alters physiological responses to fentanyl.** Stress input drives $\mathrm{PVN}^{\mathrm{CRH}}$ activity, CRH stimulates ACTH release, and ACTH promotes adrenal corticosterone production. Corticosterone provides negative feedback onto upstream CRH and ACTH signaling. The model incorporates adrenal functional mass (A) as a slower state variable, whereby $dA/dt$ is regulated by ACTH-dependent adrenal cell-mass production and adrenal cell-mass removal. All model parameters are detailed in Tables S3.

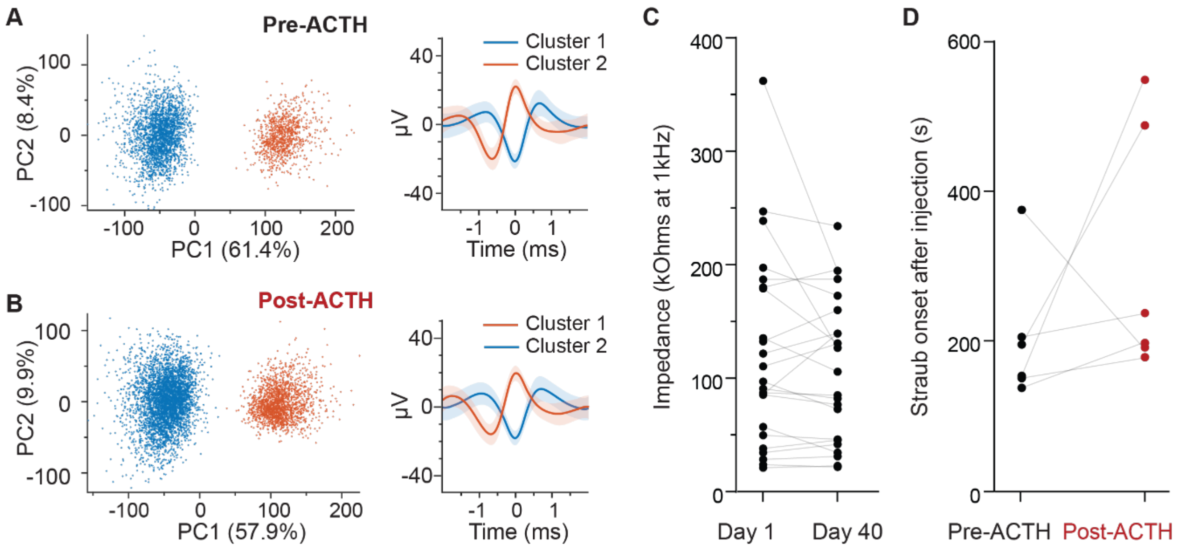


**Fig. S9 Electrophysiology recording quality remains stable over the course of the study. (A)** Baseline spike clustering by PCA of a single electrode on day 1. Average waveform ± SEM of each cluster shown. **(B) Left:** Spike clustering by PCA of the same electrode on day 40. **Right:** Lines and shaded areas represent average waveform ± standard error of the mean (SEM) of each cluster shown. **(C)** Electrode impedance (kOhms) measured at 1 kHz remained stable between recording sessions Individual electrodes are paired from day 1 (mean ± SD: 121.4 ± 85.86 kOhms) to day 40 (107.8 ± 63.68 kOhms; n = 23 electrodes across 3 mice). **(D)** Time between fentanyl injection and Straub response day 1 (mean ± SD: 202.5 ± 88.67 s) to day 40 (306.7 ± 166.4 s; paired t test: p = 0.275; difference between days mean ± SEM: 104.2 ± 85.01 s; n = 6 mice).

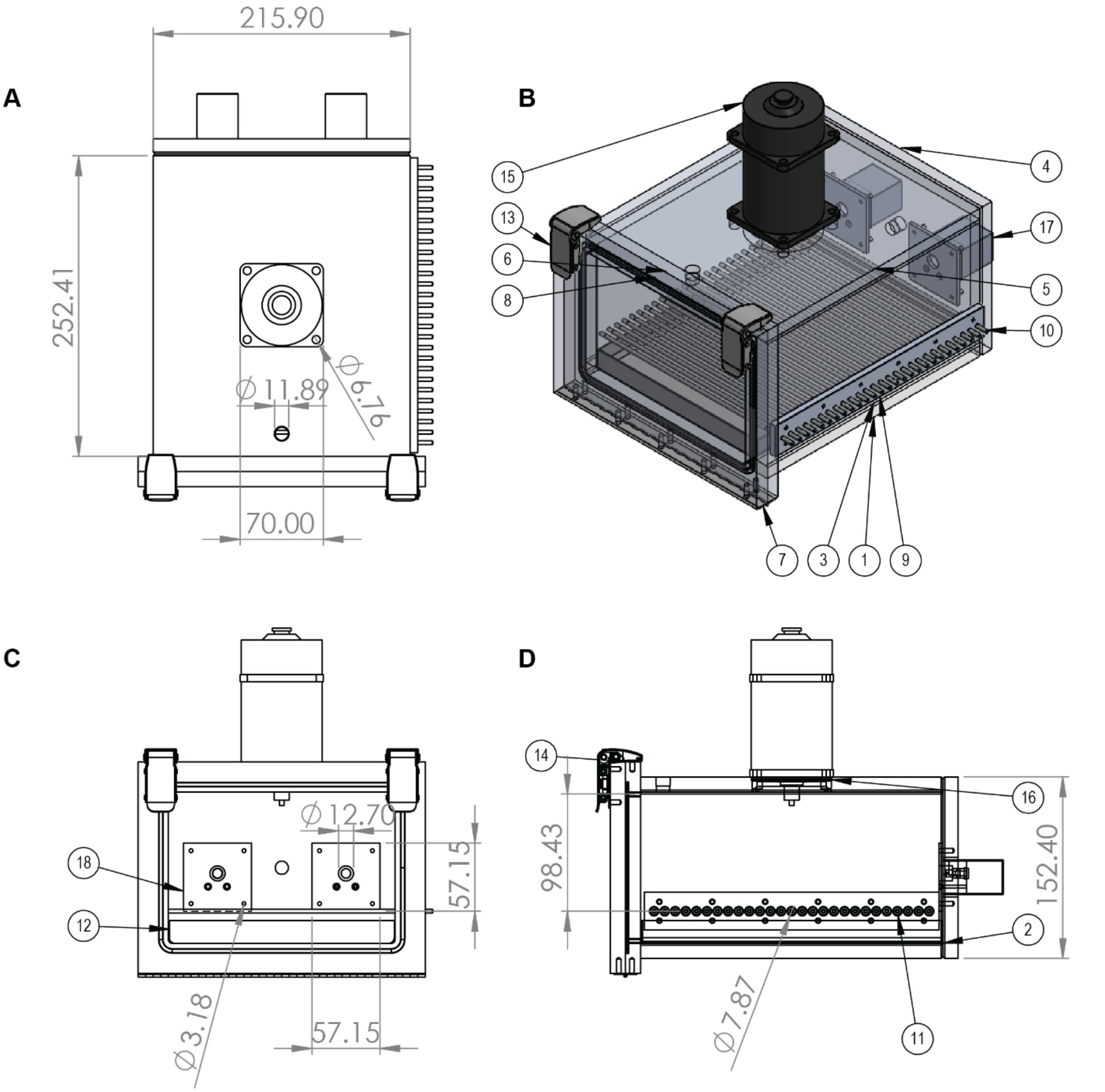


**Fig. S10 Custom operant vapor chamber enables non-surgical fentanyl self-administration.** **(A)** Top-down view showing the two nose poke devices (top) and the latched front door (bottom). **(B)** Isometric 3D assembly view of the complete chamber with commutator, and foot shock grid floor. **(C)** Front view through the latched door showing the internal chamber layout with the foot shock grid and collection tray positioned at the bottom for waste management. **(D)** Side elevation view illustrating the mechanical integration of the foot shock grid, and overall chamber profile. All units are in millimeters. Material specifications are outlined in table S4.

**Supplementary Tables**

| Display label | Full GO term | GO ID | Gene ratio | FDR adjusted p-value |
|---|---|---|---|---|
| Lipid homeostasis | regulation of lipid metabolic process | GO:0019216 | 0.064 | < 0.05 |
| Peptidase homeostasis | regulation of peptidase activity | GO:0052547 | 0.064 | < 0.05 |
| Neurogenesis regulation | regulation of neurogenesis | GO:0050767 | 0.060 | < 0.05 |
| Chemical stress response | cellular response to chemical stress | GO:0062197 | 0.050 | < 0.05 |
| Steroid metabolism | steroid metabolic process | GO:0008202 | 0.050 | < 0.05 |
| Ketone metabolism | ketone metabolic process | GO:0042180 | 0.046 | < 0.05 |
| Lipid biosynthesis regulation | regulation of lipid biosynthetic process | GO:0046890 | 0.041 | < 0.05 |
| Oxidative stress response | cellular response to oxidative stress | GO:0034599 | 0.041 | < 0.05 |
| Hormone metabolism | hormone metabolic process | GO:0042445 | 0.041 | < 0.05 |
| Collagen metabolism | collagen metabolic process | GO:0032963 | 0.037 | < 0.05 |
| Steroid biosynthesis | steroid biosynthetic process | GO:0006694 | 0.037 | < 0.05 |
| Temperature homeostasis | temperature homeostasis | GO:0001659 | 0.037 | < 0.05 |
| Amine metabolism | amine metabolic process | GO:0009308 | 0.032 | < 0.05 |
| Response to ROS | cellular response to reactive oxygen species | GO:0034614 | 0.032 | < 0.05 |
| Insulin response regulation | regulation of cellular response to insulin stimulus | GO:1900076 | 0.028 | < 0.05 |
| Collagen biosynthesis | regulation of collagen biosynthetic process | GO:0032965 | 0.025 | < 0.05 |
| Collagen homeostasis | regulation of collagen metabolic process | GO:0010712 | 0.025 | < 0.05 |
| Cell migration | substrate-dependent cell migration | GO:0006929 | 0.018 | < 0.05 |
| Metabolite generation | generation of precursor metabolites and energy | GO:0006091 | 0.132 | < 0.001 |
| Fatty acid metabolism | fatty acid metabolic process | GO:0006631 | 0.119 | < 0.001 |
| Organic compound oxidation | energy derivation by oxidation of organic compounds | GO:0015980 | 0.102 | < 0.001 |

| Display label | Full GO term | GO ID | Gene ratio | FDR adjusted p-value |
|---|---|---|---|---|
| Nucleotide metabolism | nucleotide metabolic process | GO:0009117 | 0.100 | < 0.001 |
| Purine metabolism | purine nucleotide metabolic process | GO:0006163 | 0.098 | < 0.001 |
| Small molecule catabolism | small molecule catabolic process | GO:0044282 | 0.096 | < 0.001 |
| Lipid catabolism | lipid catabolic process | GO:0016042 | 0.091 | < 0.001 |
| Cellular respiration | cellular respiration | GO:0045333 | 0.082 | < 0.001 |
| Ribonucleotide metabolism | ribonucleotide metabolic process | GO:0009259 | 0.081 | < 0.001 |
| Organic acid catabolism | organic acid catabolic process | GO:0016054 | 0.079 | < 0.001 |
| Carboxylic acid catabolism | carboxylic acid catabolic process | GO:0046395 | 0.079 | < 0.001 |
| Purine ribonucleotide metabolism | purine ribonucleotide metabolic process | GO:0009150 | 0.079 | < 0.001 |
| Aerobic respiration | aerobic respiration | GO:0009060 | 0.071 | < 0.001 |
| Cellular lipid catabolism | cellular lipid catabolic process | GO:0044242 | 0.071 | < 0.001 |
| Fatty acid oxidation | fatty acid oxidation | GO:0019395 | 0.057 | < 0.001 |
| Lipid oxidation | lipid oxidation | GO:0034440 | 0.057 | < 0.001 |
| Fatty acid catabolism | fatty acid catabolic process | GO:0009062 | 0.054 | < 0.001 |
| Fatty acid beta oxidation | fatty acid beta-oxidation | GO:0006635 | 0.047 | < 0.001 |

**Table S1. Enriched GO BP terms associated with fentanyl administration**. GO BP terms enriched among differentially expressed genes in the zona fasciculata population of fentanyl-treated relative to saline-treated adrenal glands are shown. This table provides detailed results accompanying Fig. S5, with upregulated pathways highlighted in green and downregulated pathways highlighted in blue. Gene ratios represent the fraction of input genes associated with each GO term. FDR-adjusted p-values are reported as thresholded ranges based on enrichment analysis significance thresholds (upregulated terms significant at FDR < 0.05; downregulated terms significant at FDR < 0.001). Display labels summarize corresponding full GO term descriptions.

| Treatment | Duration | Dose | Route of administration | Product, manufacturer |
|---|---|---|---|---|
| Crinecerfont | 3 days | 10 mg/kg/day | Intraperitoneal | HY-106203A, MedChem Express |
| ACTH | 4 days | 0.45 mg/kg/day | Intraperitoneal | A0298, Sigma-Aldrich |
| Saline | 4 days | 200 µL | Intraperitoneal | Sodium Chloride, Preservative Free 0.9%, Hospira |
| Fentanyl | 4 days | 0.3 mg/kg/day | Intraperitoneal | 1269902, Sigma-Aldrich |
| Naloxone hydrochloride | 4 days | 5 mg/kg/day | Intraperitoneal | BP548, Sigma-Aldrich |
| Naloxone methiodide | 4 days | 50 mg/kg/day | Intraperitoneal | N129, Sigma-Aldrich |

**Table S2 Pharmacological treatments and dosing parameters.** Pharmacological treatments and dosing parameters. All compounds were administered via intraperitoneal injection. Crinecerfont was given for 3 days; all other compounds (ACTH, fentanyl, naloxone hydrochloride, naloxone methiodide, and saline) were given for 4 days. Combinatorial treatments used the same individual parameters in sequential administration. Dose, duration, catalog numbers, and manufacturers are listed for each compound.

| Variable | Definition | Function |
| --- | --- | --- |
| CRH | Corticotropin-releasing hormone concentration | Hypothalamic signal to the pituitary |
| ACTH | Adrenocorticotropic hormone concentration | Pituitary hormone that stimulates the adrenal gland |
| CORT | Corticosterone concentration | Adrenal glucocorticoid output that feeds back onto the HPA axis |
| A | Adrenal functional mass | Slow-changing adrenal state variable that determines corticosterone-producing capacity |
| s | Stress input | External input that stimulates CRH production |
| $g_1$(CORT) | Corticosterone-dependent feedback on CRH production | Reduces CRH when corticosterone is high |
| $g_2$(CORT) | Corticosterone-dependent feedback on ACTH production | Reduces ACTH when corticosterone is high |
| $b_1$ | CRH production rate | Stress-induced CRH production |
| $a_1$ | CRH removal rate | The rate at which CRH decays or is cleared |
| $b_2$ | ACTH production rate | CRH-driven ACTH production |
| $a_2$ | ACTH removal rate | The rate at which ACTH decays or is cleared |
| $b_3$ | Corticosterone production rate | ACTH- and adrenal mass-dependent corticosterone production |
| $a_3$ | Corticosterone removal rate | Determines the rate at which corticosterone decays or is cleared |
| $b_A$ | ACTH-dependent adrenal growth coefficient | Scales the trophic effect of ACTH on adrenal functional mass |
| $a_A$ | Adrenal functional mass removal rate | Represents loss or turnover of adrenal functional mass |
| dCRH/dt | Rate of change of CRH over time | Net balance between CRH production and removal |
| dACTH/dt | Rate of change of ACTH over time | Net balance between ACTH production and removal |

| | | |
|---|---|---|
| dCORT/dt | Rate of change of corticosterone over time | Net balance between adrenal corticosterone production and removal |
| dA/dt | Rate of change of adrenal functional mass over time | Net balance between ACTH-driven adrenal growth and adrenal mass loss |

**Table S3 Model variables and parameters used to describe HPA axis dynamics.** Variables, parameters, feedback functions, rate constants, and differential equations used in the dynamic HPA-axis model. Variables include CRH, ACTH, and CORT representing fast hormone dynamics at the hypothalamic, pituitary, and adrenal levels, respectively. Adrenal functional mass (A) represents a slow-changing state variable that determines corticosterone-producing capacity. Corticosterone-dependent feedback functions (g1 and g2) regulate CRH and ACTH production. Rate constants (b and a terms) describe hormone production and clearance, while parameters bA and aA govern ACTH-dependent adrenal growth and mass loss. Differential equations (dCRH/dt, dACTH/dt, dCORT/dt, dA/dt) describe the dynamics of each state variable.

| ITEM NO. | PART NUMBER | DESCRIPTION | QTY. |
|---|---|---|---|
| 1 | BASE | Clear Acrylic | 1 |
| 2 | SIDEWALL LEFT | Clear Acrylic | 1 |
| 3 | SIDEWALL RIGHT | Clear Acrylic | 1 |
| 4 | BACKWALL | Clear Acrylic | 1 |
| 5 | TOP | Clear Acrylic | 1 |
| 6 | FRONT | Clear Acrylic | 1 |
| 7 | 12075A21 | Piano Hinge with Holes | 2 |
| 8 | DOOR | Clear Acrylic | 1 |
| 9 | ROD COVER | Clear Acrylic | 1 |
| 10 | ROD | Aluminum | 27 |
| 11 | 2418T112 | Oil-Resistant Soft Buna-N O-Rings | 27 |
| 12 | TRAY | Aluminum | 1 |
| 13 | 4432N122_Corner-Mount Draw Latch 1 | Corner-Mount Draw Latch | 2 |
| 14 | 4432N122_Corner-Mount Draw Latch 2 | Corner-Mount Draw Latch | 2 |
| 15 | Doric ARJ | Assisted Electrical Rotary Joint | 1 |
| 16 | 1173N403 | High-Temperature Soft Silicone O-Ring | 1 |
| 17 | ENV-313M | Illuminated Nose Poke | 2 |
| 18 | FACEPLATE | Black Acrylic | 2 |

**Table S4 Self-administration chamber components list.** Bill of materials detailing the structural components, mechanical hardware, electronic components, and quantities required for chamber assembly.
